# The roles of space and food web complexity in mediating ecological recovery

**DOI:** 10.1101/2025.05.13.653715

**Authors:** Klementyna A. Gawecka, Matthew A. Barbour, James M. Bullock, Jordi Bascompte

## Abstract

Landscape-scale ecological restoration is a key strategy for halting and reversing biodiversity decline. However, ensuring the long-term sustainability of restoration efforts requires guiding the recovery of complex ecological systems with many interdependent species at a landscape scale. Due to these challenges, our understanding of recovery trajectories remains limited. Using experiments and metapopulation models, we explore how the spatial configuration of communities and food web complexity jointly influence species recovery at different spatial scales. We find that the number and spatial placement of communities affects the colonisation of empty habitat patches, but does not influence population recovery in patches where communities are introduced. Food web complexity reduces the recovery of lower trophic levels, both at population and metapopulation scales. However, this negative effect may be partially mitigated at higher levels of food web complexity. Our results demonstrate that the joint consideration of spatial configuration and species interactions could enhance the effectiveness of restoration actions.

## Introduction

To ‘bend the curve’ of biodiversity loss, we must restore our degraded or lost natural habitats (Leclère et al., 2020). To ensure the persistence of species, restoration activities need to focus on landscapes – i.e. metapopulations and metacommunities – rather than communities at individual sites (Isaac et al., 2018, Bullock et al., 2002). In particular, to promote functioning ecosystems, ecological communities of multiple interacting species must be restored (Oliver et al., 2015, Tylianakis et al., 2010). Together, spatial and community complexity can promote long-term sustainability and resilience (Bullock et al., 2022).

Yet, as systems become more extensive and complex, predicting their dynamics becomes increasingly difficult. Ecological recovery is often assessed at single locations and at a limited number of timepoints by measuring species abundance or richness (e.g., Escobar et al., 2025, Hordijk et al., 2024, Banin et al., 2023, Resch et al., 2022). Success is then determined by comparing these measures to a chosen reference state (Atkinson et al., 2022). However, complex systems can by highly dynamic and follow nonlinear trajectories (Sutheimer et al., 2025). Importantly, these trajectories can vary across space, depending on the local biotic (e.g., species abundances) and abiotic (e.g., habitat type or position within the landscape) conditions. Moreover, the synchrony, or lack thereof, between local dynamics can lead to very different patterns at larger spatial scales (Firkowski et al., 2022, Wilcox et al., 2017, Laliberté et al., 2013). Understanding these spatially-structured recovery trajectories and their drivers is vital for guiding restoration actions (Montoya, 2021, Suding, 2011), and hence for effective restoration planning.

One key factor shaping recovery is spatial configuration. The arrangement and connectivity of habitat patches govern species dispersal and colonisation processes. Thus, the spatial structure of the landscape influences local community dynamics (Bowler and Benton, 2009, Bowler and Benton, 2005), species spread across the landscape (Rayfield et al., 2023, Gawecka and Bascompte, 2023, Saade et al., 2022, Gilarranz et al., 2017), and metacommunity structure (Bertellotti et al., 2023), stability (Arancibia, 2024) and persistence (Li et al., 2023, Arancibia and Morin, 2022, Gilarranz and Bascompte, 2012). Despite its clear significance, a critical question remains: how does spatial configuration affect recovery trajectories?

Restoration practice often focuses on promoting a target species or enhancing species diversity (Pettorelli and Bullock, 2023, Brudvig, 2011). However, species in a community are interdependent, through direct or indirect interactions. Species interactions influence community and metacommunity dynamics (May and Hassell, 1981, Bastolla et al., 2009), stability (Firkowski et al., 2022, Rohr et al., 2014, Thébault and Fontaine, 2010) and persistence (Domínguez-Garcia et al., 2024, Gaiarsa and Bascompte, 2022). The recovery of one species can have profound effects on others (Gawecka and Bascompte, 2021, Horn et al., 2020, Baker et al., 2019), and this impact depends on the community and landscape context (Twining et al., 2022). Yet, species interactions are rarely considered in restoration practice (Hallett et al., 2023). If restoration’s goal is to rebuild ecological complexity, we must understand how species interactions affect recovery trajectories across landscapes.

Here, we combine experimental and modelling approaches to study the landscape-scale recovery of species embedded in communities. It is difficult to examine such processes robustly in field settings, so controlled experiments combined with modelling can help elucidate mechanisms (e.g., Gilarranz et al., 2017). Specifically, we investigate how recovery trajectories are affected by (1) spatial configuration – the number and location of introduced communities, and (2) food web complexity. We assess recovery at both local (habitat patch) and global (landscape) scales. First, we conduct an experiment using model landscapes composed of five connected patches, and three insect communities varying in food web complexity. Second, using a parameterised metacommunity model, we assess the robustness of our experimental findings by simulating recovery dynamics in larger landscapes and communities.

## Methods

### Experimental design

We used three experimental communities consisting of a plant (radish, *Raphanus sativus*), two aphid species – cabbage aphid *Brevicoryne brassicae* (BB) and turnip aphid *Lipaphis erysimi* (LE) –, and a parasitoid wasp *Diaeretiella rapae* (DR) (Figure 1A). The communities increase in food web complexity: from a single aphid species – plant interaction, through the addition of interspecific competition between two aphid species, to the inclusion of parasitism. We refer to these communities as *BB, BB-LE* and *BB-LE-DR*, respectively. We focus on the response of aphid BB as it is the weaker competitor of the two species (Barbour et al., 2022), and thus its recovery may be more uncertain. We present the response of LE in the Supporting Information (SI).

**Figure 1:**
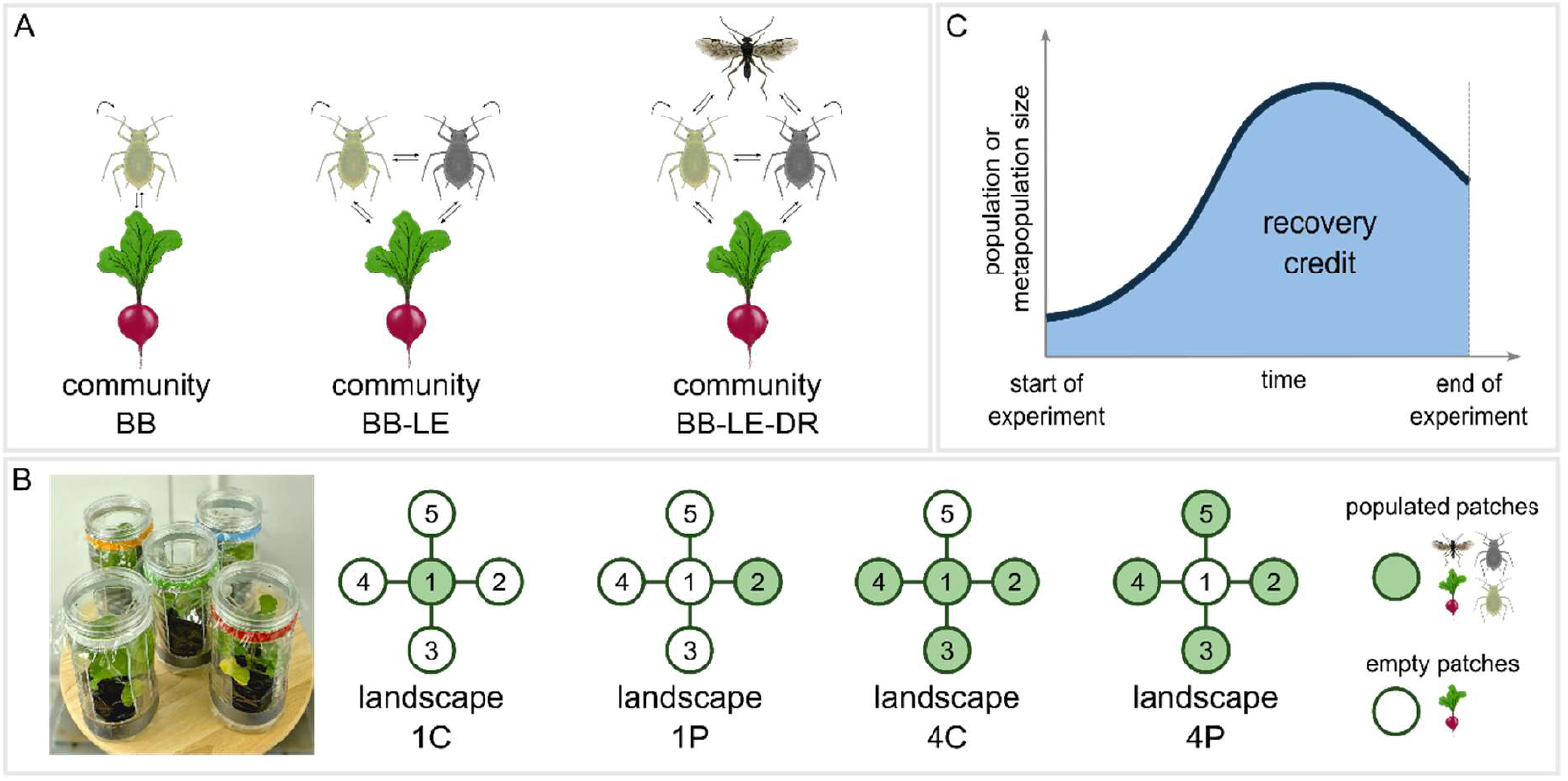
Experimental design. (A) Three experimental communities increasing in food web complexity (from left to right). The communities are composed of a plant, *Raphanus sativus*, aphids *Brevicoryne brassicae* (BB) and *Lipaphis erysimi* (LE), and a parasitoid wasp, *Diaeretiella rapae* (DR).(B)Experimental landscapes consisting of five containers (habitat patches) connected in a star configuration. Initially, only selected patches contain a plant and an insect community (*populated* patches, green on the schematic). Other patches initially contain only a plant (*empty* patches, white on the schematic). (C) We quantify recovery trajectory as recovery credit – the area under the curve of the population or metapopulation size against time, for the duration of the experiment (shaded area).

Our experimental landscape comp five habitat patches in a star configuration with one central patch and four peripheral patches (Figure 1B), designed to allow assessment of the effects of spatial configuration of introduced communities without confounding by varying overall landscape structure. A habitat patch was represented by a cylindrical, transparent polyethylene container (diameter 10 cm, height 20 cm). There were four openings on the side and one in the lid, which were covered with transparent cellophane to allow air exchange. The bottom of the container had holes for water drainage and was covered with nylon mesh to prevent insects escaping. The containers housed a 7 cm x 7 cm x 6 cm plant pot. The patches were connected with a silicone tube (diameter 1 cm) at the top of the container. A string running through the tube further enabled the insects to migrate between the patches.

Initially, all five patches contained a single, two-week-old plant. However, the landscapes differed in how many, and in which patches we placed an insect community (one of the three communities). We considered four landscapes with an insect community initially in: one central patch (landscape *1C*), one peripheral patch (landscape *1P*), four central patches (landscape *4C*), and four peripheral patches (landscape *4C*) (Figure 1B). We refer to the patches containing a community as *populated*, and patches with initially only a plant as *empty*.

We applied these community and landscape treatments in a fully factorial design, resulting in 12 unique combinations. We replicated each combination five times.

### Experimental procedure

We seeded plants two weeks prior to the start of the experiment with two seeds per pot. After one week, we reduced the number to a single plant per pot. The plants were grown in a greenhouse and watered once per week.

We reared aphids and parasitoid wasps in mesh cages in a climate chamber set to 22 °C, 50% humidity and 16 h photoperiod. Aphids sourced for the experiment were maintained on the same radish species as used in the experiment. The parasitoid wasps were reared on a non-experimental aphid species (green peach aphid, *Myzus persicae*). For more details on the insect colonies, refer to Barbour et al. (2022).

At the start of the experiment, we placed ten aphids of each species on a two-week-old plant inside the relevant containers (i.e., *populated* patches, Figure 1B). In the case of community *BB-LE-DR*, we transferred a single one-day-old, mated female parasitoid wasp into the same containers as the aphids. Finally, we placed the experimental units on trays in the climate chamber (set to 22 °C, 50% humidity and 16 h photoperiod) for the duration of the experiment. We positioned the units in random orientations and locations within the climate chamber and shuffled every week.

We counted aphids of each species in each container twice a week (every 3 or 4 days, see Figure S 2-Figure S 3 for aphid count time series). We watered the experimental units once a week by filling trays with water such that the bottoms of plant pots were just submerged. The experiment ran for 26 days, which covered approximately four generations of aphids, two generations of parasitoid wasps and the lifespan of the plants.

### Metacommunity model

We developed a spatially explicit model based on the mass-effect paradigm of metacommunity theory (Leibold et al., 2004). It describes the local dynamics of our communities and dispersal across the landscape. Its general form resembles other discrete-time mass-effect models with Lotka-Voleterra-type competition and predation (e.g., Thompson and Gonzalez, 2017). However, we chose the specific functions such that the model reproduced the observed dynamics of our experimental communities (see SI). This approach balances the model’s generality and precision (Levins, 1966). The population size of aphid species *i* in patch *k* at time *t* is given by:

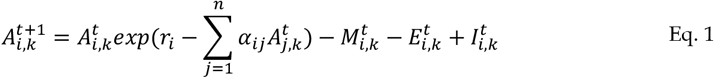

The first term describes density-dependent intrinsic growth, intraspecific competition, and interspecific competition with aphid species *j. r*_*i*_ is the intrinsic growth rate, *α*_*ij*_ is the intraspecific (when *j* = *i*) or interspecific competition coefficient (when *j* ≠ *i*), and *n* is the number of aphid species. We found that an exponential form of the logistic growth model (e.g., Agrawal, 2004) best describes the dynamics of our experimental system. The second term represents the mortality due to parasitism and depends on the density of the parasitoid. We adopted a piecewise linear saturating function which mimics a Type II functional response (Eq. S 3). The final two terms describe density-dependent emigration from patch *k* (Eq. S 4) and immigration into patch *k* from adjacent patches (Eq. S 5), respectively. The parasitoid population size was modelled as a balance between births, deaths, emigration, and emigration (Eq. S 6 – Eq. S 8). We provide details of all model functions in the SI.

We parameterised the model using a series of experiments, each designed to enable the determination of a parameter(s). For example, to estimate the intrinsic growth rate and intraspecific competition of aphid BB, we (1) experimentally measured its population size in a single patch over time, (2) fitted a linear model to per capita growth rate (in terms of log-transformed differences) versus its population size, and (3) obtained confidence intervals of the intercept (intrinsic growth rate) and slope (intraspecific competition). We carried out these parameterisation experiments under the same conditions, and using the same plant species and insect colonies, as in the main experiment. We provide more details on the parameterisation procedure in the SI.

To validate the parameterised model, we simulated the main experiment and compared the model output to the experimental observations. We found that, despite its relative simplicity and generality, our model qualitatively reproduces the experimental trends (Figure S 7-Figure S 12). Importantly, this allows us to use the model to disentangle various processes otherwise masked in the experiment, as well as test robustness of experimental findings by simulating alternative scenarios.

First, we considered larger landscapes and alternative configurations. We simulated the dynamics of the experimental communities in two landscapes with 13 habitat patches (see insets in Figure 3). In the *spread* configuration, the patches were placed in a star with four arms, whereas in the *clustered* configuration, there were 12 patches directly connected to a central patch. As in the experiment, we introduced either one or four communities into either the most central or most peripheral patches.

Second, we considered two more complex food webs. One community consisted of three aphid species and the parasitoid wasp (community *BB-LE-A3-DR*), whereas the other included two aphids, the parasitoid wasp and a hyperparasitoid wasp (community *BB-LE-DR-HP*). The hyperparasitoid wasp lays eggs inside the parasitoid’s larvae, thus reducing the number of emerged parasitoids. For illustrative purpose, all model parameters of the new aphid species were assumed to be the averages of the corresponding BB’s and LE’s parameters. In the case of the hyperparasitoid wasp, we adopted the same dynamics and model parameters as those of the parasitoid.

### Recovery measure

To quantify the recovery trajectory, we computed the recovery credit (Marjakangas et al., 2018, Hanski, 2000) which is analogous to the recovery debt (Moreno-Mateos et al., 2017), but represents the surplus in population or metapopulation size. We defined our recovery credit as the area under the curve of the population or metapopulation size against time (shaded area in Figure 1C). In other words, our recovery credit is an integrative measure of (meta)population size through time. We calculated this credit for each aphid species, and at two spatial scales. At the local scale, we considered the population size in each patch, differentiating between initially *populated* and *empty* patches (Figure 1B). This allowed us to study the local dynamics within introduced populations, and colonisation of the rest of the landscape. At the global scale, we evaluated the recovery of metapopulation size – the sum of population sizes across all patches, which represents the overall species recovery across a landscape.

### Statistical analysis

We performed three-way ANOVA to assess the effects of (1) the number, (2) the location, and (3) the food web complexity of the introduced communities on the recovery credit. We carried out separate tests for the recovery credit of (a) populations in *empty* patches, (b) populations in *populated* patches, and (c) metapopulations of each aphid species. In the case of local population recovery, we considered the average recovery credit across all initially *empty* or *populated* patches in each landscape.

To ensure normality and include zero values of recovery credit, we applied ln(*x* + 1) transformation to the calculated recovery credit (for untransformed results, see Figure S 4-Figure S 5). We report the results in terms of average effect as a percentage change, and provide the ANOVA tables in SI (Table S 2-Table S 7). We performed all statistical analyses and model simulations in R version 4.4.0 (R Core Team, 2020).

## Results

### Experiment

We find that recovery in initially *empty* patches is affected by the number and location of introduced communities, as well as food web complexity (Figure 2A). Recovery credit is 26% greater in landscapes where communities are introduced into four patches compared to one (*F*_*1,50*_ = 10.8, *P* = 0.0019), and 21% greater where the introductions are into peripheral compared to central patches (*F*_*1,50*_ = 6.7, *P* = 0.012). However, we find a significant interaction between the number and location of communities (*F*_*1,50*_ = 6.2, *P* = 0.016). Post-hoc comparisons reveal that the effect of the number of initial communities is substantial only in landscapes where communities are introduced into peripheral patches (landscapes *1P* and *4P*). In turn, community location has an effect only in landscapes with four introduced communities (landscapes *4C* and *4P*). Overall, food web complexity reduces recovery (*F*_*2,50*_ = 5.4, *P* = 0.0072). While the addition of aphid LE does not appear to affect BB’s recovery (1.3% reduction in recovery credit, comparing communities *BB* and *BB-LE*), the parasitoid reduces the aphid’s recovery credit by 22% (comparing communities *BB-LE* and *BB-LE-DR*). Aphid LE shows a qualitatively similar response (Figure S 6A), except for the effect of community location which is not statistically significant.

**Figure 2:**
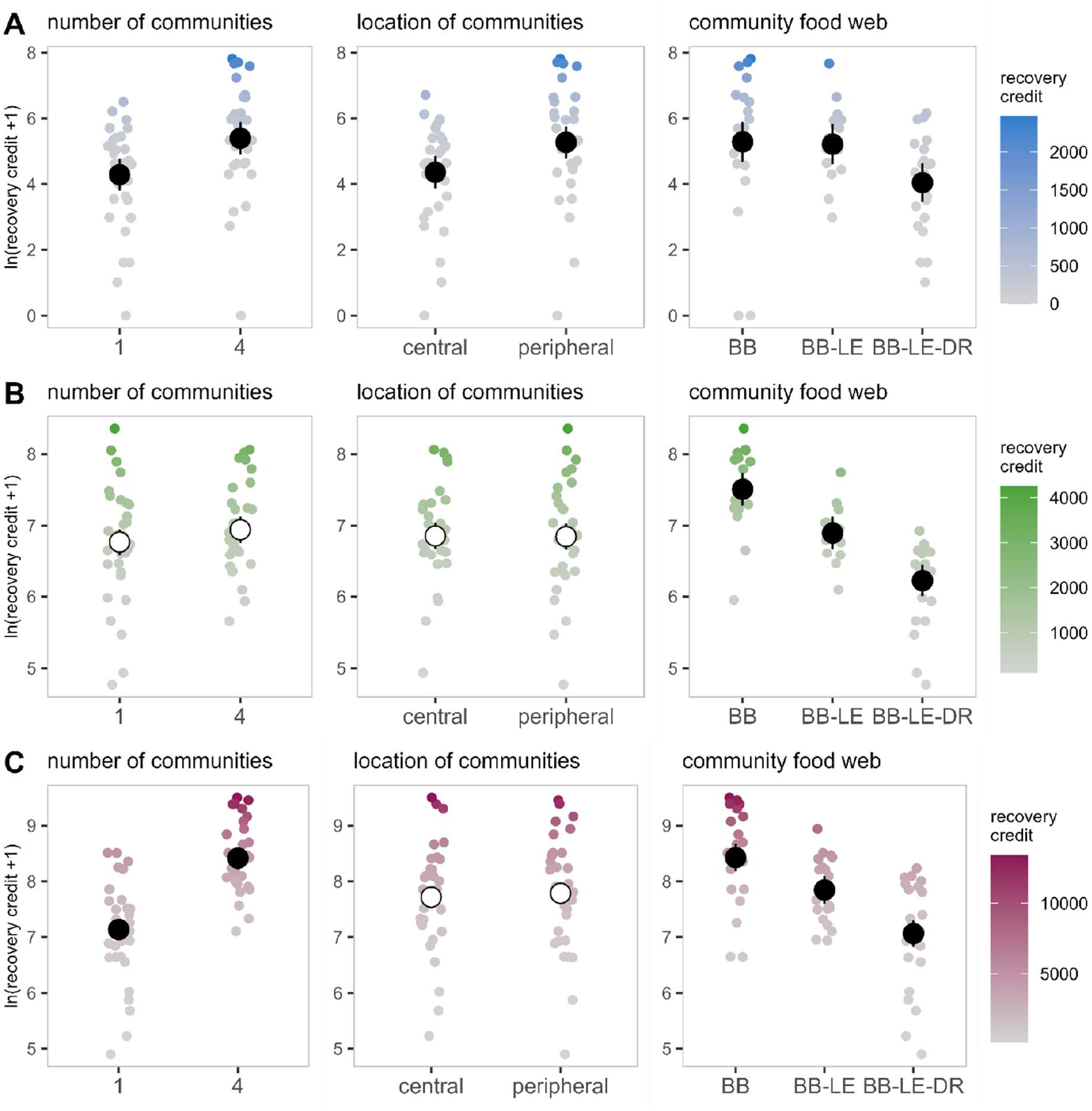
Recovery of aphid BB (A) in initially *empty* patches, (B) in initially *populated* patches, and (C)metapopulation. Panels show the effects of initial number of communities, location of initial communities, and community food web on the recovery credit (*ln*(*x* + 1) transformed). Smaller points represent recovery credit calculated from experimental data, with colours indicating their untransformed values. Larger points and vertical lines depict average linear model predictions and their 95% confidence intervals, respectively. Full and empty points indicate statistically significant and nonsignificant effects, respectively.

Conversely, the recovery in initially *populated* patches is not influenced by the spatial configuration (the number or location of initial communities, Figure 2B). However, increasing food web complexity substantially reduces the aphid’s recovery (*F*_*2,50*_ = 32.5, *P* < 0.001, Figure 2B). The addition of aphid LE and the parasitoid decreases BB’s recovery by 8% (comparing communities *BB* and *BB-LE*), and 10% (comparing communities *BB-LE* and *BB-LE-DR*), respectively. Aphid LE shows a qualitatively similar response (Figure S 6B).

At the *metapopulation* scale, recovery is affected by the number of introduced communities and food web complexity, but there is no clear effect of their initial location (Figure 2C). Increasing the number of initial communities from 1 to 4 boosts the recovery credit by 18% (*F*_*1,50*_ = 87.7, *P* < 0.001), whereas increasing food web complexity reduces it (*F*_*2,50*_ = 32.1, *P* < 0.001). Specifically, the addition of aphid LE decreases BB’s recovery credit by 7% (comparing communities *BB* and *BB-LE*), whereas the parasitoid causes a further reduction of 10% (comparing communities *BB-LE* and *BB-LE-DR*). We find no clear effect of interactions between the initial number of communities and their complexity. Aphid LE shows a qualitatively similar response (Figure S 6C).

### Simulated scenarios

Our model simulations indicate that the relationship between the number and location of introduced communities and recovery in initially *empty* patches may be landscape-dependent (Figure 3). For example, the *spread* patch configuration, where 13 patches are arranged in a four-armed star, has higher recovery credit when four communities are introduced compared to only one. However, the differences between central and peripheral introductions are not substantial (Figure S 14). In the *clustered* configuration, with 12 peripheral patches connected to a single central patch, recovery credit is lowest in *1P* landscape, whereas the differences between landscapes *1C, 4C* and *4P* are negligible. Similar to the experiment, recovery credit in the initially *populated* patches appears to be affected only by the food web complexity (Figure S 14 B, Figure S 15 B). *Metapopulation* recovery credit is influenced by both the number and complexity of introduced communities, as in the experimental scenario (Figure S 14 C, Figure S 15 C).

**Figure 3:**
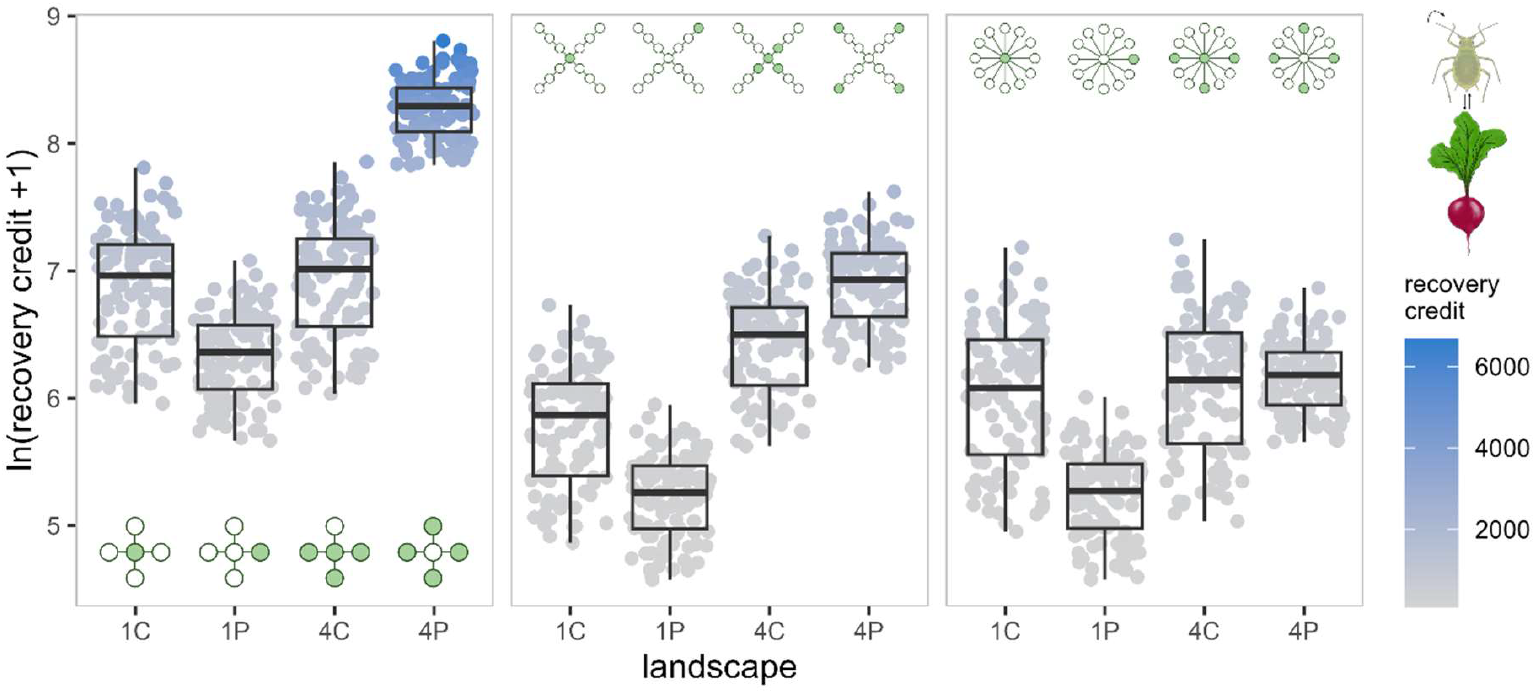
Simulated recovery of aphid BB in *BB* community in initially *empty* patches. Panels corresponds to various simulated patch configurations: *experimental* (left), *spread* (middle) and *clustered* (right). Points plot the recovery credit (*ln*(*x* + 1) transformed) calculated from model simulations. Landscapes *1C, 1P, 4C* and *4P* indicate the number and location of introduced communities.

We find that, in the model, the addition of a third aphid species to the *BB-LE-DR* community increases the recovery credit of aphid BB at both local and global scales (Figure 4). Moreover, the presence of a hyperparasitoid increases BB’s recovery further, relative to community *BB-LE-DR*. The effects of the number and location of initial communities in these simulations are qualitatively similar to those observed in the experiment (Figure S 16).

**Figure 4:**
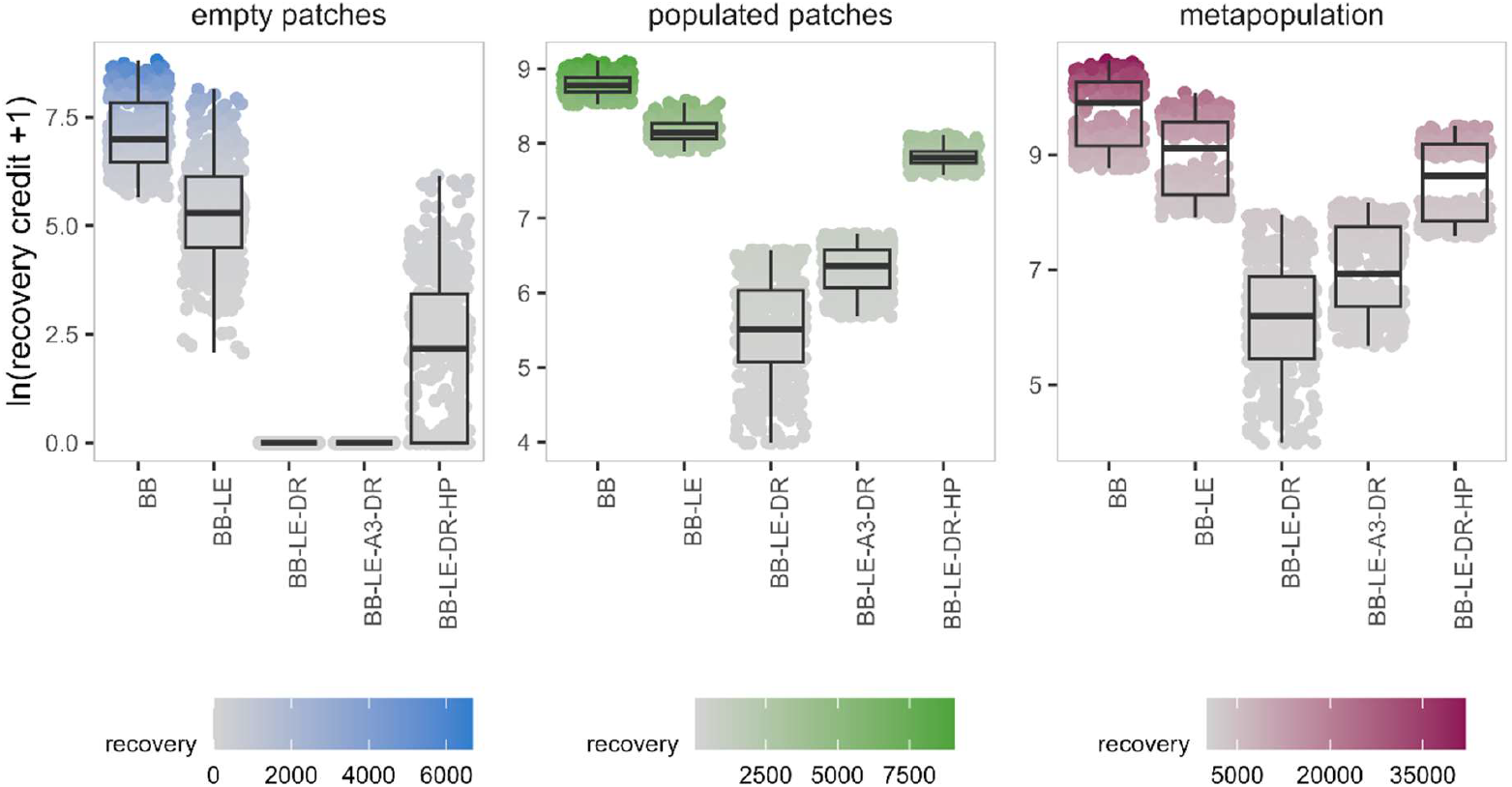
Simulated recovery of aphid BB in various communities in the experimental landscape configuration. Panels corresponds to recovery in *empty* patches (left), in *populated* patches (middle) and of *metapopulation* (right). Points plot the recovery credit (*ln*(*x* + 1) transformed) calculated from model simulations. Boxplots summarise all landscapes (*1C, 1P, 4C* and *4P*). A3 refers to a third aphid species, and HP to a hyperparasitoid wasp.

## Discussion

We find that both spatial configuration – the number and location of introduced communities –, and food web complexity affect species recovery. Yet, the importance of these factors depends on the process of interest – local population growth or landscape colonisation. Both processes are key to ecological restoration, and our analysis suggests important consequences for nature recovery actions.

### The role of spatial configuration

Generally, increasing the number of introduced communities increases recovery of the metapopulation, and of the populations colonising habitat patches which are initially empty. A greater number of communities implies more potential sources, and thus, increases the chances of an empty patch being colonised and a population establishing. Furthermore, introducing communities into peripheral, poorly connected habitat patches may be beneficial for population recovery in initially empty patches. Interestingly, peripheral patches have been shown to be important for initiating cascades across networks (Van De Leemput et al., 2024). Yet, this process may depend on the number of introduced communities and the patch arrangement within the landscape. Our experimental results suggest that prioritising peripheral patches is particularly beneficial when we introduce four communities, but may not have substantial impacts with only one initial community. In other words, introducing multiple peripheral communities boosts the colonisation of central patches. These patches, in turn, are important for linking the landscape, enabling movement across it and ensuring metapopulation persistence (e.g., Cumming et al., 2022, Thompson et al., 2017, Gilarranz et al., 2015).

An experiment by Saade et al. (2022) also found that the recolonization dynamics following patch extinctions depend the spatial distribution of the intact patches, although less so on their number. Our model simulations indicate that interaction between the number and location of introduced communities may depend on the spatial landscape structure. For example, in a landscape with many peripheral patches connected to a single central patch, colonisation of empty patches is slowest when a single peripheral community is introduced, but there is no effect of location with four introduced communities. The dependence of the spread of individuals across the landscape on patch configuration has also been demonstrated by Rayfield et al. (2023) and Gilarranz et al. (2017) using experiments and models. In summary, we find that how many and where communities are introduced affects the dispersal pathways, and thus, the colonisation of the landscape.

Yet, the number and location of introduced communities has little effect on the recovery of the populations in patches where they are introduced. Similarly, Altermatt et al. (2011) found in an experiment that local populations in undisturbed patches are unaffected by emigration of individuals into disturbed patches. This suggests that the population growth in these patches is driven by local intra- and inter-specific dynamics, rather than immigration and emigration. In fact, our metacommunity model allows us to decompose the contribution of local population processes and dispersal to the change in population size at a given time (Figure S 13). First, we find that the initially populated patches act as sources, with emigration outweighing immigration. Conversely, the initially empty patches are sinks with net immigration. Second, the contribution of dispersal to the change in population size is much smaller in the initially *populated* than *empty* patches (see also Bird et al., 2024, Bullock et al., 2020).

However, we note the timescale of our experiment is relatively short and the observed dynamics are transient. As hinted by the model results, the relative contribution of local processes and dispersal may be highly dynamic (Figure S 13). We postulate that, as local populations fluctuate, the rescue effect of immigration demonstrated by many previous studies (e.g., Liu and Vidal, 2025, Li et al., 2023, Staddon et al., 2010, Holyoak, 2000, Gonzalez et al., 1998) may become more prevalent (see Ives et al., 2004). Moreover, the contribution of dispersal has been shown to depend on dispersal rate (Zelnik et al., 2019, Thompson et al., 2017, Altermatt et al., 2011) and the dispersal kernel (Rayfield et al., 2023). This points to not only species-specificity in the effects of spatial configuration on recovery, but also the influence of the landscape matrix (e.g., Fletcher et al., 2024, Aström and Pärt, 2013).

### The role of food web complexity

The complexity of the introduced communities has a profound effect on recovery at both local and global scales. The recovery of our focal aphid species reduces upon the addition of another aphid species (i.e., interspecific competition), and the parasitoid wasp (i.e., parasitism). By impeding population growth locally, these interspecific interactions reduce the overall dispersal potential and recovery at the landscape scale. However, there is empirical evidence that interspecific interactions affect dispersal, and thus potentially colonisation, in other ways. For example, Fronhofer et al. (2018) found that both top-down and bottom-up control increased active emigration rates. Allbee et al. (2023) found that herbivores influenced dispersal distances of plants. And a review by Bestion et al. (2024) concluded that detrimental interactions increased dispersal, whereas beneficial ones reduced it. While it is possible that these mechanisms are at work in our system, we observe that they are swamped by the negative effects of competition and parasitism on population growth (see also Bullock et al., 2020).

However, the negative effects of interspecific competition and parasitism could be, at least partially, offset by even greater food web complexity. For example, our model simulations suggest that the addition of another aphid species increases the focal aphid’s recovery relative to the community with two aphid species and a parasitoid wasp. This positive effect, despite another source of interspecific competition, is due to reduced parasitism rate on each aphid species, i.e., a dilution effect (Foster and Treherne, 1981). Alternatively, the addition of a higher trophic level species increases aphid’s recovery by reducing the parasitism pressure, i.e., mesopredator suppression (Ritchie and Johnson, 2009, May and Hassell, 1981) (but see Horn, 1989 for evidence of more complex spatial dynamics at play). In summary, higher food web complexity allows for more indirect effects among species, affecting recovery in less predictable ways.

### Future directions

Here, we study ecological recovery from the perspective of a herbivore. Yet, different species and species guilds may be affected by space and community differently, and thus follow different recovery trajectories. This is perhaps most obvious in species involved in trophic interactions where recovery of the resource benefits the consumer, but not vice versa (as shown here, also see Gawecka and Bascompte, 2021). However, even among species belonging to the same guild, recovery depends on the number of interaction partners (Gawecka and Bascompte, 2023). Additionally, species dispersal abilities can vary widely across trophic levels (Elzinga et al., 2007). This affects how species perceive the landscape (e.g., Bertellotti et al., 2023), and thus, the role of spatial configuration in recovery. In short, recovery trajectories of species at various trophic levels in species- and interaction-rich communities remains to be investigated.

Our landscapes consist of equally sized patches with identical habitat. However, both habitat area and type influence species distributions. Moreover, these effects can be species-specific (e.g., Dong et al., 2025, Gardner et al., 2024, Twining et al., 2022, Van Noordwijk et al., 2015), and may influence interspecific interactions (Lennox et al., 2025). While our experimental approach found no interaction between spatial configuration (number and location of communities) and food web complexity, we postulate that such interaction may become important in more heterogeneous landscapes.

## Conclusions

There is a trade-off between species recovery and food web complexity. Yet, we need complexity for ecosystem functioning and resilience (Liang et al., 2025, Tilman et al., 2014). We propose three approaches to this conundrum:

1. spatial planning which considers landscape structure – prioritising many peripheral patches (or a single central patch if resources are limited) for introductions may aid landscape colonisation,
2. building species- and interaction-rich communities – indirect effects within communities may boost the recovery relative to species-poor communities,
3. staggered species introductions – allowing lower trophic levels to establish before introducing higher trophic levels.

Our results suggest that ecological restoration involves a delicate balancing act. Despite this, much restoration practice takes little account of community or spatial complexities (Maes et al., 2024, Bullock et al., 2022). However, by integrating species interactions and spatial landscape configuration into restoration planning, it is likely that we can enhance recovery.

## Acknowledgements

We thank Daniel Trujillo-Villegas for maintaining the insect colonies, and assistance constructing and trialling the experimental setup, as well as conducting the experiment. We also thank Marcel Freund for help constructing the experimental setup, Matthias Furler for greenhouse support, and Tobias Züst and Kyle Coblentz for advice on the experimental design. Access to an additional climate chamber was kindly provided by Rie Shimizu-Inatsugi and Kentaro Shimizu.

## Funding

This work was supported by the University of Zurich Postdoc Grant (grant FK-22-114) and Marie Skłodowska-Curie Actions Postdoctoral Fellowship (grant EP/Z000831/1) to KAG, Natural Sciences and Engineering Research Council of Canada (NSERC) Discovery Grant and Fonds de recherche du Québec–Nature (FRQNT) Relève Professorale Grant to MAB, Natural Environment Research Council (NERC) consortium award ‘Restoring Resilient Ecosystems’ (grant NE/V006525/1) to JMB, and Schweizerischer Nationalfonds zur Förderung der Wissenschaftlichen Forschung (SNSF, grant 310030_197201) to JB.

## Data availability

All data and code used in this study is available on GitHub (github.com/kgawecka/recovery_foodweb_experiment).

## Notes

https://github.com/kgawecka/recovery_foodweb_experiment

### Competing Interest Statement

The authors have declared no competing interest.

