## Supporting Information for "The roles of space and food web complexity in mediating ecological recovery"

### Metacommunity model

#### Description

We developed a metacommunity dynamics model to help us disentangle various processes taking place in the experimental landscape. To facilitate comparison with experimental data and parameterisation, we adopted a discrete-time model, assuming a timestep of one day. We parameterised the model using data from a series of experiments performed on the insect species prior to the main experiment (see Supporting Information).

The population size of aphid species $i$ in patch $k$ at time $t$ is given by:

$$A_{i,k}^{t+1} = G_{i,k}^{t} - M_{i,k}^{t} - E_{i,k}^{t} + I_{i,k}^{t} \qquad \text{Eq. S 1}$$

The first term includes the intrinsic growth, and intra- and inter-specific competition, and is given by:

$$G_{i,k}^{t} = A_{i,k}^{t} exp(r_i - \sum_{j=1}^{n} \alpha_{ij} A_{j,k}^{t}) \qquad \text{Eq. S 2}$$

where $r_i$ is the intrinsic growth rate, $\alpha_{ij}$ is the intraspecific (when $j = i$) or interspecific competition (when $j \neq i$), and $n$ is the number of aphid species. We found that this exponential form of the logistic growth model best reproduced the experimental data.

The second term in Eq. S 1 represents mortality due to parasitism. Based on the experimental observations, we modelled it as a Type I functional response with a threshold for maximum number of parasitised aphids, $M_{max}$ (thus mimicking a Type II response):

$$M_{i,k}^{t} = \begin{cases} \beta_i A_{i,k}^{t} f P_k^t & if \quad \sum_{i=1}^{2} M_{i,k}^{t} \leq \frac{M_{max}}{\lambda} P_k^t \\ c_i \frac{M_{max}}{\lambda} P_k^t & otherwise \end{cases} \qquad \text{Eq. S 3}$$

where $\beta_i$ is the parasitoid attack rate on aphid $i$, and $f$ is the fraction of female parasitoids in a population, $P_k^{t-1}$ is the parasitoid population size, and $\lambda$ is parasitoid's lifespan (days). $M_{max}$

represents the total number of parasitised aphids throughout parasitoid's lifetime. Although the parasitisation rate has been shown to reduce throughout parasitoid's lifetime (Soni and Kumar, 2021), we assume a constant rate for simplicity. If this per-timestep quota is exceeded, we weigh the number of parasitised aphids of each species by $c_i = \beta_i A_{i,k}^{t-1} f P_k^{t-1} / \sum_{i=1}^{n} M_{i,k}^{t}$. This ensures that the differences between aphids, in terms of their abundances and parasitoid attack rates, are accounted for.

The third term in Eq. S 1 models the density-dependent aphid emigration from patch $k$ and is given by:

$$E_{i,k}^{t} = \begin{cases} 0 & if \quad A_{i,k}^{t} < A_{min,i} \\ e_i(A_{i,k}^{t} - A_{min,i}) & if \quad A_{i,k}^{t} \geq A_{min,i} \end{cases} \qquad \text{Eq. S 4}$$

where $A_{min,i}$ is the minimum population size for emigration and $e_i$ is the emigration rate.

The final term in Eq. S 1 represents immigration of aphids into patch $k$ from adjacent patches and id given by:

$$I_{i,k}^{t} = \sum_{l=1}^{m} \frac{E_{i,l}^{t}}{c_l} \qquad \text{Eq. S 5}$$

where $m$ is the number of directly connected patches to patch $k$, and $c_l$ is the number of their direct connections.

We modelled the population size of the parasitoid in patch $k$ as:

$$P_k^{t+1} = P_k^t + B_k^t - D_k^t - E_k^t + I_k^t \qquad \text{Eq. S 6}$$

where emigration ($E_k^t$) and immigration ($I_k^t$) expressions have the same form as those for aphids (Eq. S 4 and Eq. S 5). $B_k^t$ is the number of births defined as:

$$B_k^t = \begin{cases} 0 & if \quad t \leq \tau \\ \sum_{i=1}^{n} M_{i,k}^{t-\tau} & if \quad t > \tau \end{cases} \qquad \text{Eq. S 7}$$

where $\tau$ is the time between parasitisation of an aphid and emergence of an adult parasitoid, and $M_{i,k}^{t-\tau}$ (Eq. S 3) is the number of aphids parasitised at time $t - \tau$.

The number of parasitoid deaths, $D_k^t$, is given by:

$$D_k^t = \begin{cases} P_k^t & if \quad t = \lambda \\ \dfrac{P_k^t}{\sum_{k=1}^{n} P_k^t} \displaystyle\sum_{k=1}^{m} B_k^{t-\lambda} & if \quad t > \tau + \lambda \\ 0 & otherwise \end{cases} \qquad \text{Eq. S 8}$$

The first expression ensures that initially placed parasitoid die at time $t = \lambda$. The second expression represents deaths of the emerged parasitoids. It weighs the total number of deaths across all patches ($\sum_{k=1}^{m} B_k^{t-\lambda}$) by the number of parasitoids in patch $k$ relative to the total number of parasitoids. This accounts for movement of individuals between patches.

#### Parameterisation

We conducted all parameterisation experiments in the same climate chamber (22 °C, 50 % humidity and 16 h photoperiod) and using the same containers (Figure 1B) as in the main experiment. We used the same radish plant species, aphid and parasitoid colonies as in the main experiment. In all cases, we used a single, two-week old radish plants per pot. When selecting aphid individuals, we ensured that all are adults of similar size (and age). The parasitoid wasps were maintained on a non-experimental aphid species (green peach aphid, *Myzus persicae*), and supplemented with honey solution throughout experiments.

**Single aphid species response experiment**

Parameters determined: $r_i$, $\alpha_{ii}$, $e_i$, $A_{min,i}$

Experimental procedure: We connected two patches and placed a radish plant in each patch. We placed 5 aphids of the same species (BB or LE) onto the radish leaves in one of the patches (patch 1). We ran the experiment for 28 days. Three days per week (Mondays, Wednesdays and Fridays), we counted aphids in both patches, and then removed all aphids present in the initially empty patch (patch 2). This allowed us to determine the population growth in patch 1, and emigration into patch 2. We replicated this setup five times per aphid species.

Parameter determination: We determined the per capita growth rate, accounting for emigrated aphids, $E_i^{t-1}$, and number of days between counts, $\Delta t$. We plotted it against population size ($A_i^{t-1}$) and fitted a linear model. We found that the linear model fitted the experimental data best when we defined the per capita growth rate as: $\frac{\ln(A_i^t+E_i^{t-1})-\ln(A_i^{t-1})}{\Delta t}$ (Agrawal, 2004). This results in the exponential expression given by Eq. S 2. We extracted the 95% confidence intervals of the intercept (i.e., $r_i$) and slope (i.e. $\alpha_{ij}$). The minimum population size for emigration, $A_{min,i}$, was the aphid count in patch 1 when we found aphids in patch 2 for the first time. For each aphid species, we took the average value across all replicas. To determine the emigration rate, we plotted the number of emigrated aphids per day against aphid count in patch 1, and fitted a linear model with $A_{min,i}$ as the x-axis intercept. We then obtained the 95% confidence interval of the slope as $e_i$.

**Two aphid species response experiment**

Parameters determined: $\alpha_{ij}$

Experimental procedure: In a single container, we placed a radish plant and transferred 5 BB and 5 LE aphids. We ran the experiment for 28 days, counting aphids three times per week (Mondays, Wednesdays, Fridays). We replicated the experiment five times.

Parameter determination: To determine interspecific competition, we plotted $r_i - \alpha_{ii}A_i^{t-1} - \frac{\ln(A_i^t+E_i^{t-1})-\ln(A_i^{t-1})}{\Delta t}$ against $A_j^{t-1}$. For $r_i$ and $\alpha_{ii}$, we substituted the estimates determined from the single aphid species response experiments (see above). We fitted a linear model with the intercept at the origin and extracted the 95% confidence interval of the slope as $\alpha_{ij}$.

**Maximum parasitisation experiment**

Parameters determined: $M_{max}$

Experimental procedure: In a patch, we placed a radish plant and transferred 64 aphids of the same species (BB or LE) onto plant's leaves. We added a single one-day old female parasitoid. We maintained the container until mummies could be identified (i.e., beyond parasitoid's lifetime), at which point we counted them to determine the total number of parasitised aphids per parasitoid. We replicated the experiment five times per aphid species.

Parameter determination: As there was no substantial difference in the total number of parasitised aphids between the two aphid species, we determined $M_{max}$ as the average mummy count across both aphid species and all replicas.

**Parasitoid functional response experiment**

Parameters determined: $\beta_i$

Experimental procedure: We placed a radish plant in a single patch and transfer 4, 8, 16, 32 or 64 aphids of the same species (BB or LE). We added a single one-day old female parasitoid. After 24 hours, we removed the parasitoid. We maintained the patches in constant conditions until the mummies could be identified. We replicated each aphid density five times.

Parameter determination: For each aphid species, we plotted the number of mummies against the initial number of aphids. We fitted a linear model with zero intercept and obtained the 95% confidence interval of the slope as $\beta_i$.

**Time to emergence experiment**

Parameters determined: $\tau$

Experimental procedure: This experiment was a continuation of the functional response experiment. Every day, we counted the number of emerged parasitoids and removed them from the patch.

Parameter determination: We determined the time to emergence as the mean number of days since parasitism, weighted by the number of emerged parasitoids that day. Due to lack of clear difference between the two aphid species, we averaged $\tau$ across BB and LE.

**Fraction of parasitoid females experiment**

Parameters determined: $f$

Experimental procedure: We collected parasitoids from a colony of newly emerged parasitoids in batches of 5-20 individuals. We counted the number of females in each batch.

Parameter determination: We determined the fraction of females across all collected individuals.

**Parasitoid lifespan experiment**

Parameters determined: $\lambda$

Experimental procedure: We placed five female parasitoids in a patch with a radish plant. Every day, we recorded the number of dead parasitoids. We replicated this setup five times.

Parameter determination: We determined the lifespan as the mean age at death, weighted by the number of dead parasitoids found dead at that age.

**Parasitoid dispersal experiment**

Parameters determined: $e_P$, $P_{min}$

Experimental procedure: We connected two patches, each containing a single radish plant. We transferred 10 LE aphids into each patch. We added 2, 5 or 10 parasitoids into one of the patches (patch 1). We counted the number of parasitoids in both patches for three days. We replicated this experiment five times for each parasitoid density.

Parameter determination: We plotted the number of emigrated parasitoids per day (i.e., parasitoids in patch 2) against the number of parasitoids in patch 1 on the previous day. We fitted a linear model, and obtained the minimum parasitoid population size for emigration, $P_{min}$, as the x-axis intercept as and the emigration rate, $e_P$ as the slope.

As we obtained all model parameters independently of each other, we refined their estimated ranges through iterative model simulations. First, we ran 1000 model simulations which reproduced either the single aphid or two aphid species response parameterisation experiments (see above). Every iteration, we sampled model parameters from uniform distributions defined by the confidence intervals obtained. We repeated this for communities of BB only, LE only and BB & LE. Second, for every iteration, we calculated the error as the total absolute difference between the experimental observations and the corresponding model predictions. Third, we weighted each iteration by its error. We applied a linear weighting where iterations with the smallest and largest errors had weights of one and zero, respectively. Fourth, we selected 50% of iterations with the lower error. We did this independently for each community. Fifth, we obtained the interquartile range for each parameter weighted by the error in each iteration. In this final step, we considered all three communities collectively. This allowed us to determine the parameters that best reproduce both the single aphid and two aphid responses. We show model reproduction of the parameterisation experiments in Figure S 1 and the final model parameters in Table S 1.

Table S 1 Model parameter values or ranges estimated from parameterisation experiments.

| **model parameter** | **estimated value or range** | | |
|---|---|---|---|
| | **BB** | **LE** | **DR** |
| growth rate, $r_i$ | [0.314, 0.403] | [0.306, 0.397] | - |
| intraspecific competition, $\alpha_{ii}$ | [0.000697, 0.00104] | [0.000242, 0.000383] | - |
| interspecific competition, $\alpha_{ij}$ | [0.000922, 0.00115] | [0.00129, 0.00176] | - |
| parasitoid attack rate, $\beta_i$ | [0.424, 0.855] | [0.290, 0.591] | - |
| maximum number of parasitised aphids, $M_{max}$ | - | - | 62.1 |
| emigration rate, $e_i$ [day$^{-1}$] | [0.0176, 0.0257] | [0.0603, 0.101] | 0.175 |
| minimum population size for emigration, $A_{min,i}$ or $P_{min}$ | 144.4 | 167.6 | 1.5 |
| lifespan, $\lambda$ [day] | - | - | 7 |
| time to emergence, $\tau$ [day] | - | - | 16 |
| fraction of females | - | - | 0.79 |

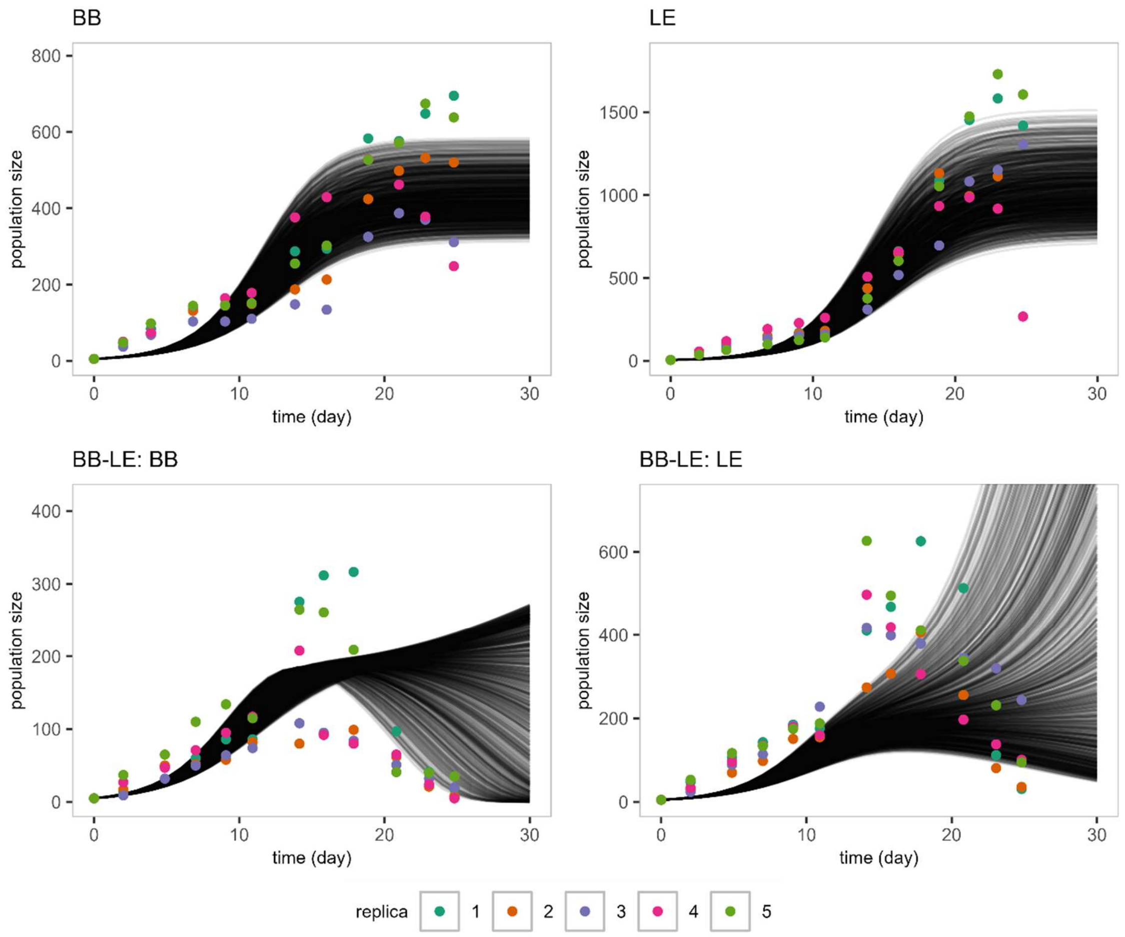


Figure S 1 Parameterisation experiments and their model simulations. Top panels show single aphid responses, whereas bottom panels depict aphid responses in two aphid communities. Points correspond to the experimental data, with colours indicating the replicas. Lines correspond to iterations of model simulations.

### Experimental results

#### Recovery trajectories

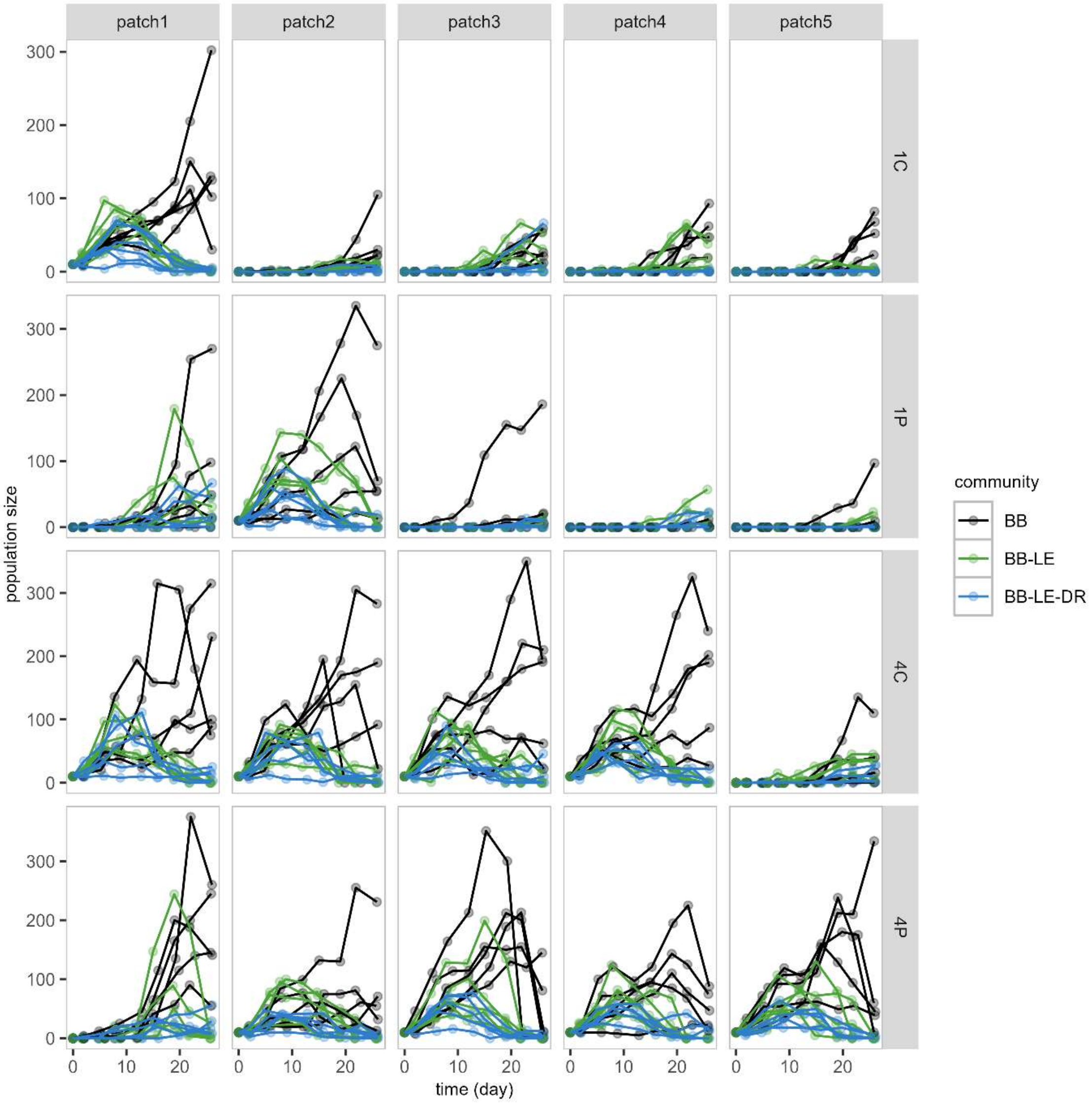


Figure S 2 Observed evolution of aphid BB population size with time. Lines join observations belonging to the same experimental replica. Colours indicate insect communities. Panels correspond to different landscapes (rows) and patches (columns).

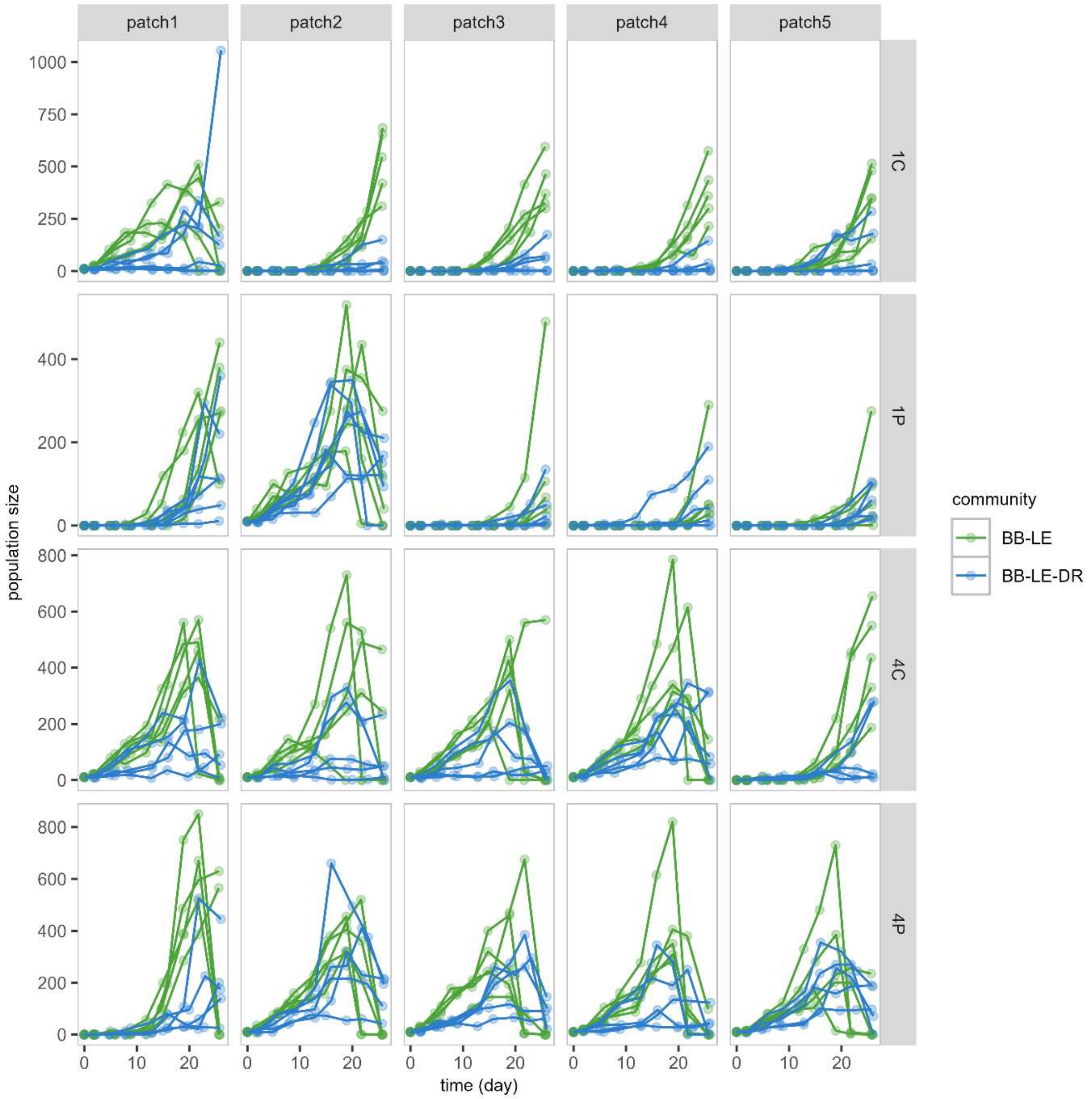

Figure S 3 Observed evolution of aphid LE population size with time. Lines join observations belonging to the same experimental replica. Colours indicate insect communities. Panels correspond to different landscapes (rows) and patches (columns).

### Recovery credit

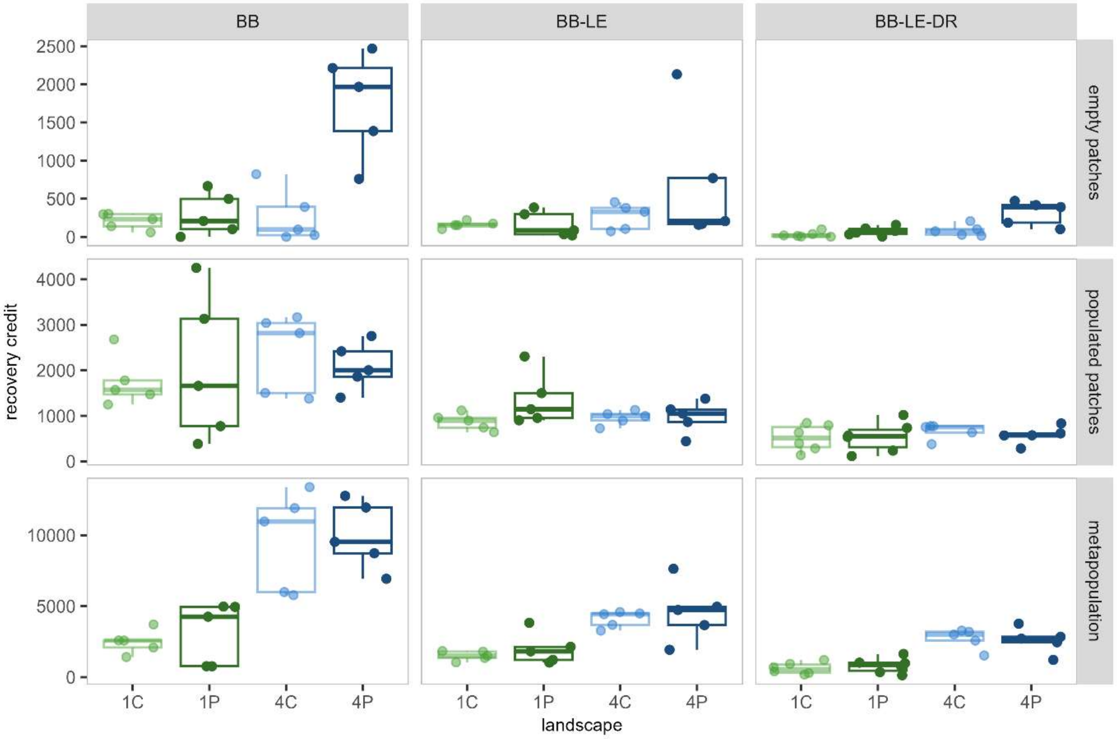


Figure S 4 Experimental recovery of aphid BB. Points plot the recovery credit calculated from experimental data. Panels correspond to different scales (rows) and communities (columns).

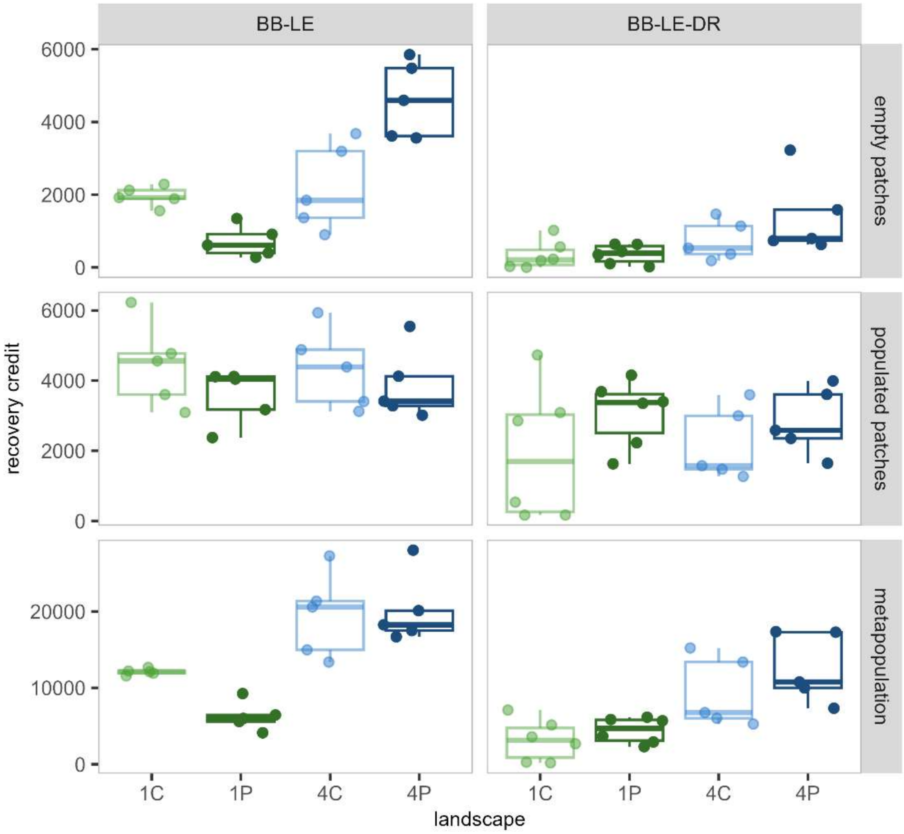


Figure S 5 Experimental recovery of aphid LE. Points plot the recovery credit calculated from experimental data. Panels correspond to different scales (rows) and communities (columns).

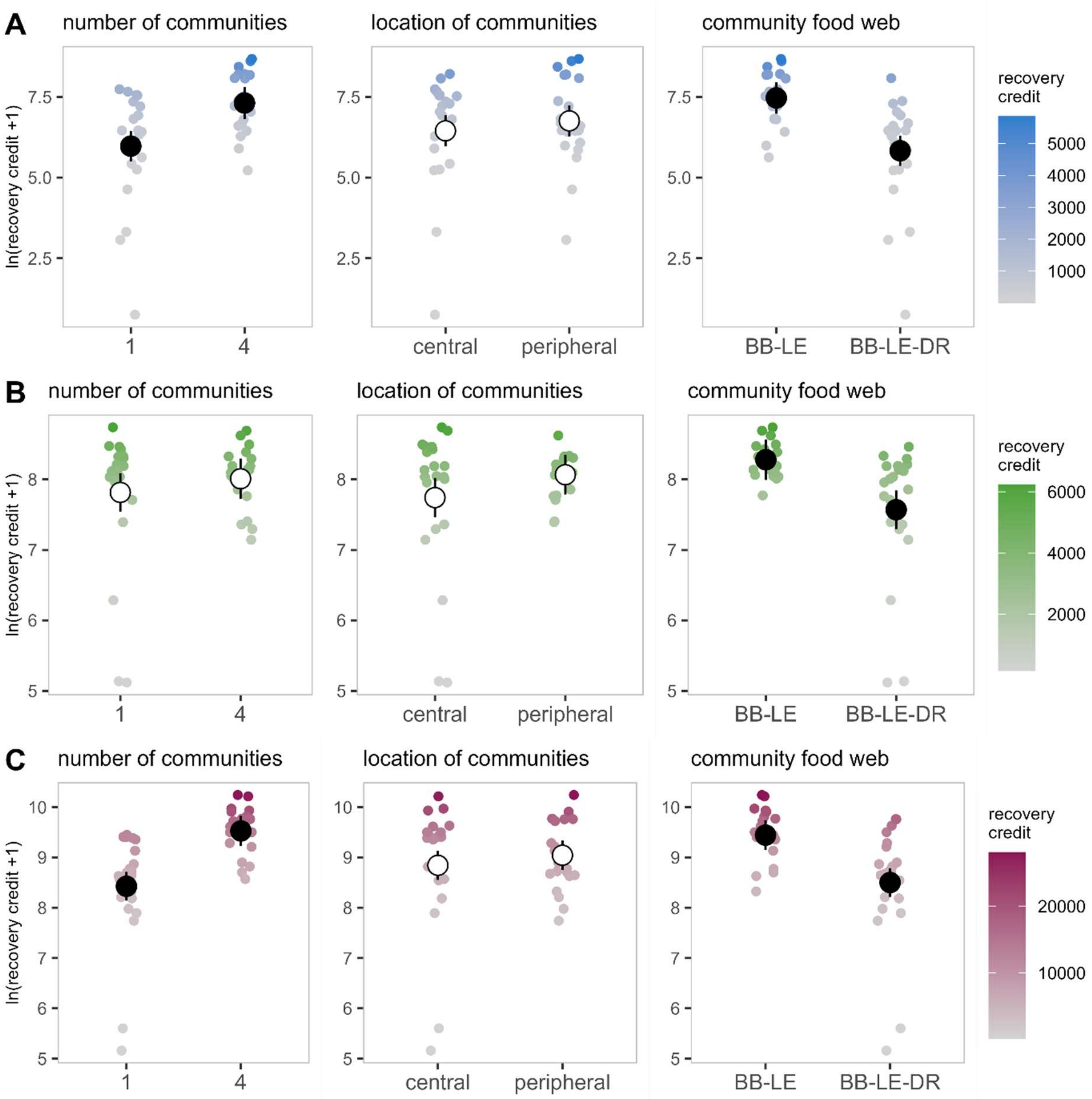


Figure S 6 Experimental recovery of aphid LE (A) in initially *empty* patches, (B) in initially *populated* patches, and (C) metapopulation. Panels show the effects of initial number of communities, location of initial communities, and community food web on the recovery credit ($ln(x+1)$ transformed). Smaller points represent recovery credit calculated from experimental data, with colours indicating their untransformed values. Larger points and vertical lines depict average linear model predictions and their 95% confidence intervals, respectively. Full and empty points indicate statistically significant and insignificant effects, respectively.

### ANOVA results

Table S 2 ANOVA of the effects for number of communities, location of communities, and community food web on the recovery credit of aphid BB in empty patches.

| | Degrees of freedom | Sum of Squares | Mean Square | F-value | P-value |
|---|---|---|---|---|---|
| number of communities | 1 | 20.752 | 20.7522 | 10.7836 | 0.001874 |
| location of communities | 1 | 12.926 | 12.9255 | 6.7166 | 0.012489 |
| community food web | 2 | 20.96 | 10.4801 | 5.4458 | 0.00725 |
| number of communities x location of communities | 1 | 11.949 | 11.9489 | 6.2091 | 0.016071 |
| location of communities x community food web | 2 | 1.473 | 0.7364 | 0.3826 | 0.684035 |
| location of communities x community food web | 2 | 6.034 | 3.0168 | 1.5676 | 0.218613 |
| number of communities x location of communities x community food web | 2 | 9.038 | 4.5189 | 2.3482 | 0.105994 |
| Residuals | 50 | 96.221 | 1.9244 | | |

Table S 3 ANOVA of the effects for number of communities, location of communities, and community food web on the recovery credit of aphid LE in empty patches.

| | Degrees of freedom | Sum of Squares | Mean Square | F-value | P-value |
|---|---|---|---|---|---|
| number of communities | 1 | 20.827 | 20.8266 | 16.5192 | 0.000269 |
| location of communities | 1 | 0.996 | 0.9957 | 0.7897 | 0.380422 |
| community food web | 1 | 27.966 | 27.9655 | 22.1817 | 4.07E-05 |
| number of communities x location of communities | 1 | 2.049 | 2.0493 | 1.6255 | 0.210977 |
| location of communities x community food web | 1 | 1.078 | 1.0777 | 0.8548 | 0.361712 |
| location of communities x community food web | 1 | 2.226 | 2.2264 | 1.766 | 0.192733 |
| number of communities x location of communities x community food web | 1 | 2.825 | 2.8246 | 2.2404 | 0.143670 |
| Residuals | 34 | 42.865 | 1.2607 | | |

Table S 4 ANOVA of the effects for number of communities, location of communities, and community food web on the recovery credit of aphid BB in populated patches.

| | Degrees of freedom | Sum of Squares | Mean Square | F-value | P-value |
|---|---|---|---|---|---|
| number of communities | 1 | 0.6986 | 0.6986 | 2.5957 | 0.1135 |
| location of communities | 1 | 0.001 | 0.001 | 0.0038 | 0.9513 |
| community food web | 2 | 17.5011 | 8.7506 | 32.5134 | 9.00e-10 |
| number of communities x location of communities | 1 | 0.1189 | 0.1189 | 0.442 | 0.5092 |
| location of communities x community food web | 2 | 0.6193 | 0.3097 | 1.1506 | 0.3247 |
| location of communities x community food web | 2 | 0.2785 | 0.1393 | 0.5174 | 0.5992 |
| number of communities x location of communities x community food web | 2 | 0.1522 | 0.0761 | 0.2827 | 0.7549 |
| Residuals | 50 | 13.4568 | 0.2691 | | |

Table S 5 ANOVA of the effects for number of communities, location of communities, and community food web on the recovery credit of aphid LE in populated patches.

| | Degrees of freedom | Sum of Squares | Mean Square | F-value | P-value |
|---|---|---|---|---|---|
| number of communities | 1 | 0.5311 | 0.5311 | 1.2441 | 0.272508 |
| location of communities | 1 | 1.095 | 1.095 | 2.5649 | 0.118509 |
| community food web | 1 | 5.2229 | 5.2229 | 12.2341 | 1.33E-03 |
| number of communities x location of communities | 1 | 0.4961 | 0.4961 | 1.1621 | 0.288624 |
| location of communities x community food web | 1 | 0.2572 | 0.2572 | 0.6025 | 0.442996 |
| location of communities x community food web | 1 | 2.1666 | 2.1666 | 5.075 | 0.030839 |
| number of communities x location of communities x community food web | 1 | 0.5902 | 0.5902 | 1.3824 | 0.247856 |
| Residuals | 34 | 14.5151 | 0.4269 | | |

Table S 6 ANOVA of the effects for number of communities, location of communities, and community food web on the recovery credit of aphid BB metapopulation.

| | Degrees of freedom | Sum of Squares | Mean Square | F-value | P-value |
|---|---|---|---|---|---|
| number of communities | 1 | 27.2292 | 27.2292 | 87.7044 | 1.39E-12 |
| location of communities | 1 | 0.0751 | 0.0751 | 0.242 | 0.6249 |
| community food web | 2 | 19.9125 | 9.9562 | 32.0687 | 1.09E-09 |
| number of communities x location of communities | 1 | 0.0507 | 0.0507 | 0.1634 | 0.6878 |
| location of communities x community food web | 2 | 0.9552 | 0.4776 | 1.5383 | 0.2247 |
| location of communities x community food web | 2 | 0.0209 | 0.0104 | 0.0336 | 0.967 |
| number of communities x location of communities x community food web | 2 | 0.0992 | 0.0496 | 0.1597 | 0.8528 |
| Residuals | 50 | 15.5233 | 0.3105 | | |

Table S 7 ANOVA of the effects for number of communities, location of communities, and community food web on the recovery credit of aphid LE metapopulation.

| | Degrees of freedom | Sum of Squares | Mean Square | F-value | P-value |
|---|---|---|---|---|---|
| number of communities | 1 | 13.6533 | 13.6533 | 29.4287 | 4.82E-06 |
| location of communities | 1 | 0.4192 | 0.4192 | 0.9037 | 0.34851 |
| community food web | 1 | 9.2566 | 9.2566 | 19.9518 | 8.35E-05 |
| number of communities x location of communities | 1 | 0.0007 | 0.0007 | 0.0014 | 0.97001 |
| location of communities x community food web | 1 | 0.7858 | 0.7858 | 1.6938 | 0.20185 |
| location of communities x community food web | 1 | 2.5999 | 2.5999 | 5.6039 | 0.02375 |
| number of communities x location of communities x community food web | 1 | 1.1879 | 1.1879 | 2.5603 | 0.11883 |
| Residuals | 34 | 15.7741 | 0.4639 | | |

### Model simulation of experiment

#### Recovery trajectories

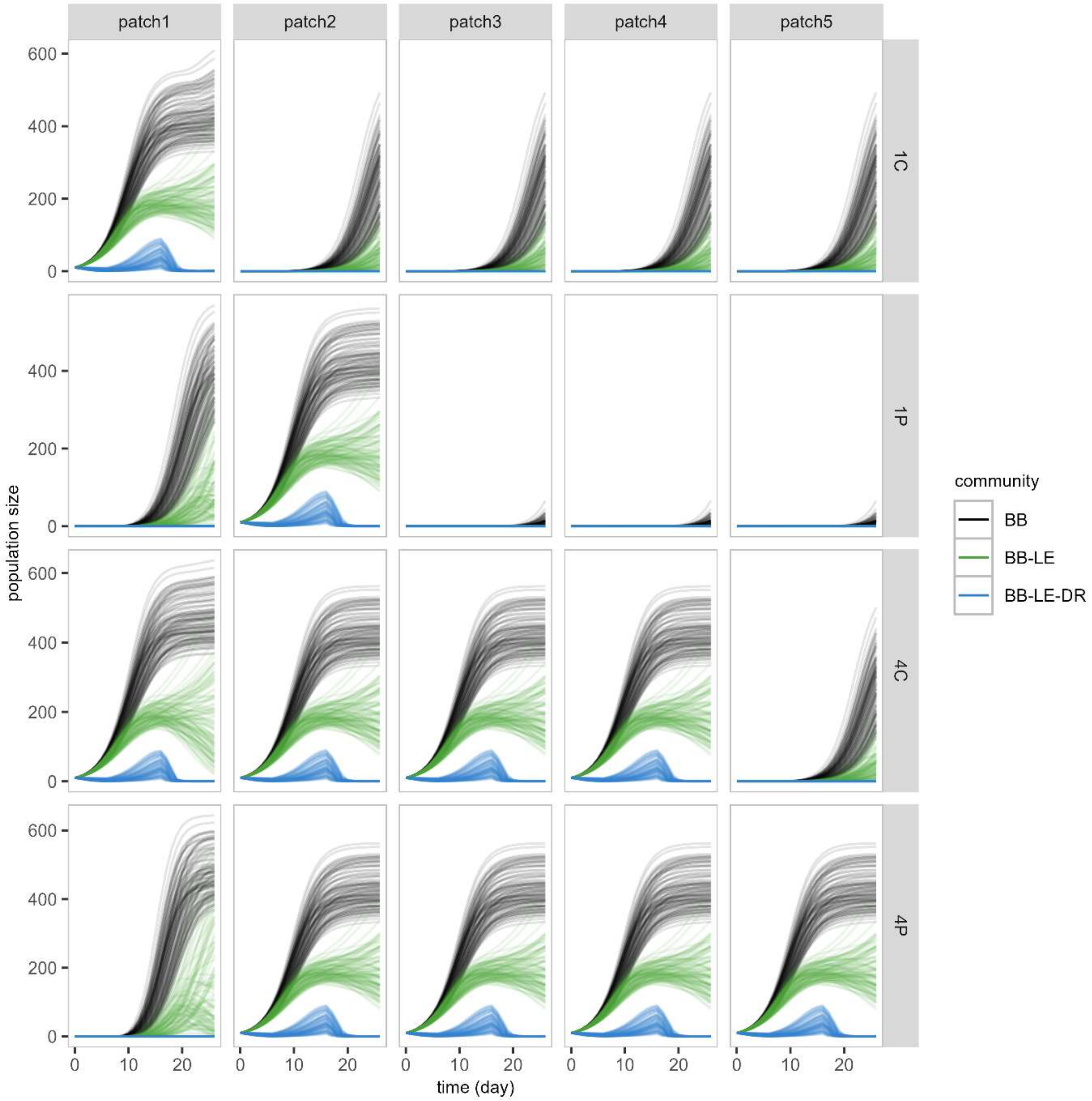


Figure S 7 Simulated evolution of aphid BB population size with time. Lines join observations belonging to the same simulation replica. Colours indicate insect communities. Panels correspond to different landscapes (rows) and patches (columns).

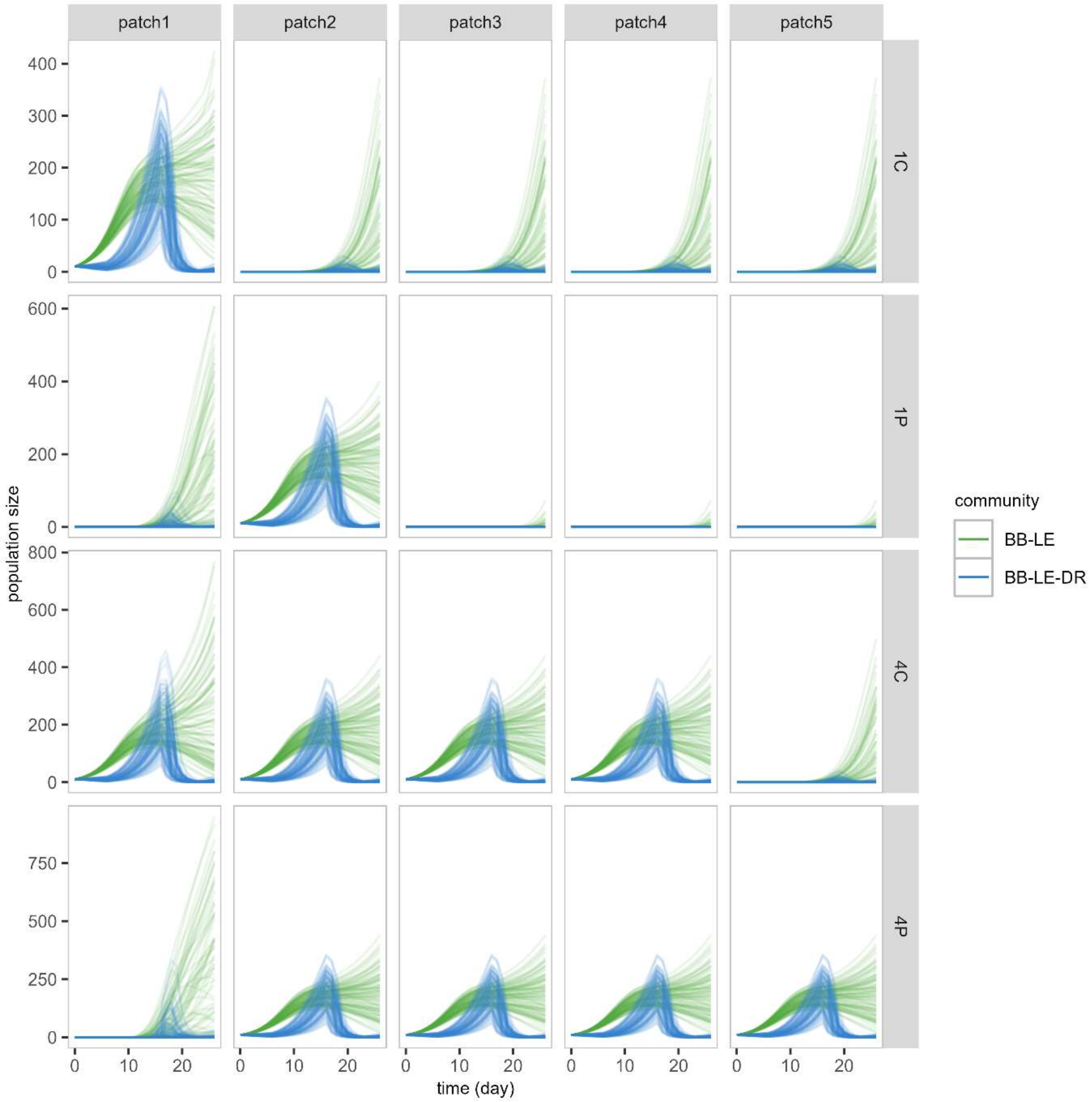

Figure S 8 Simulated evolution of aphid BB population size with time. Lines join observations belonging to the same simulation replica. Colours indicate insect communities. Panels correspond to different landscapes (rows) and patches (columns).

#### Recovery credit

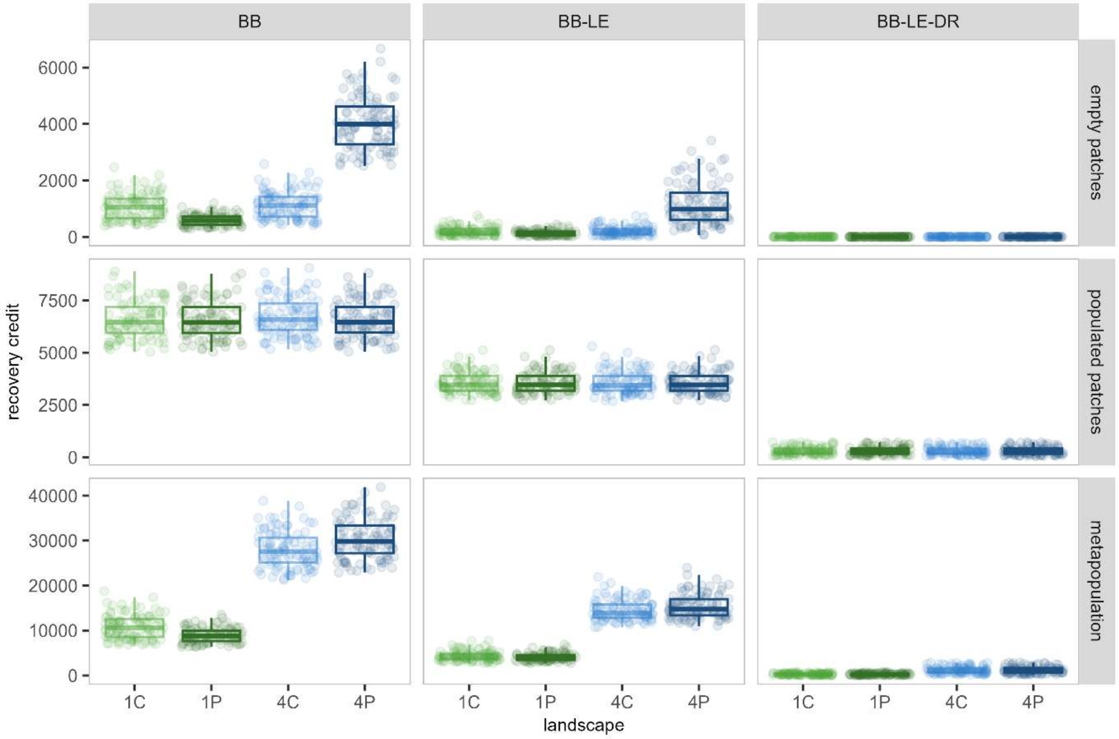


Figure S 9 Simulated recovery of aphid BB. Points plot the recovery credit calculated from model simulations. Panels correspond to different scales (rows) and communities (columns).

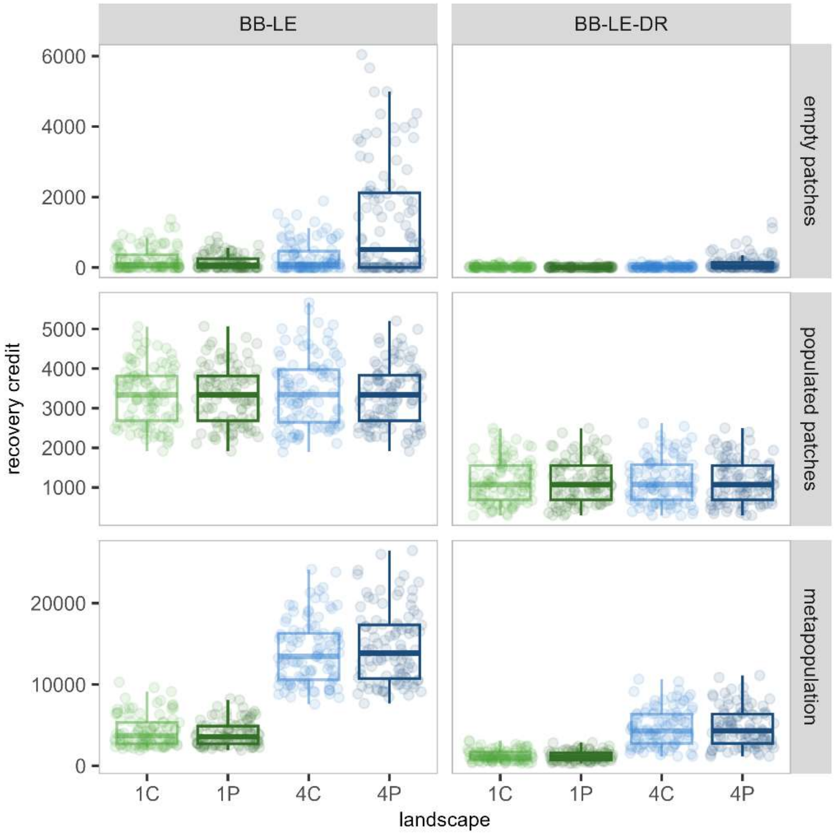


Figure S 10 Simulated recovery of aphid LE. Points plot the recovery credit calculated from model simulations. Panels correspond to different scales (rows) and communities (columns).

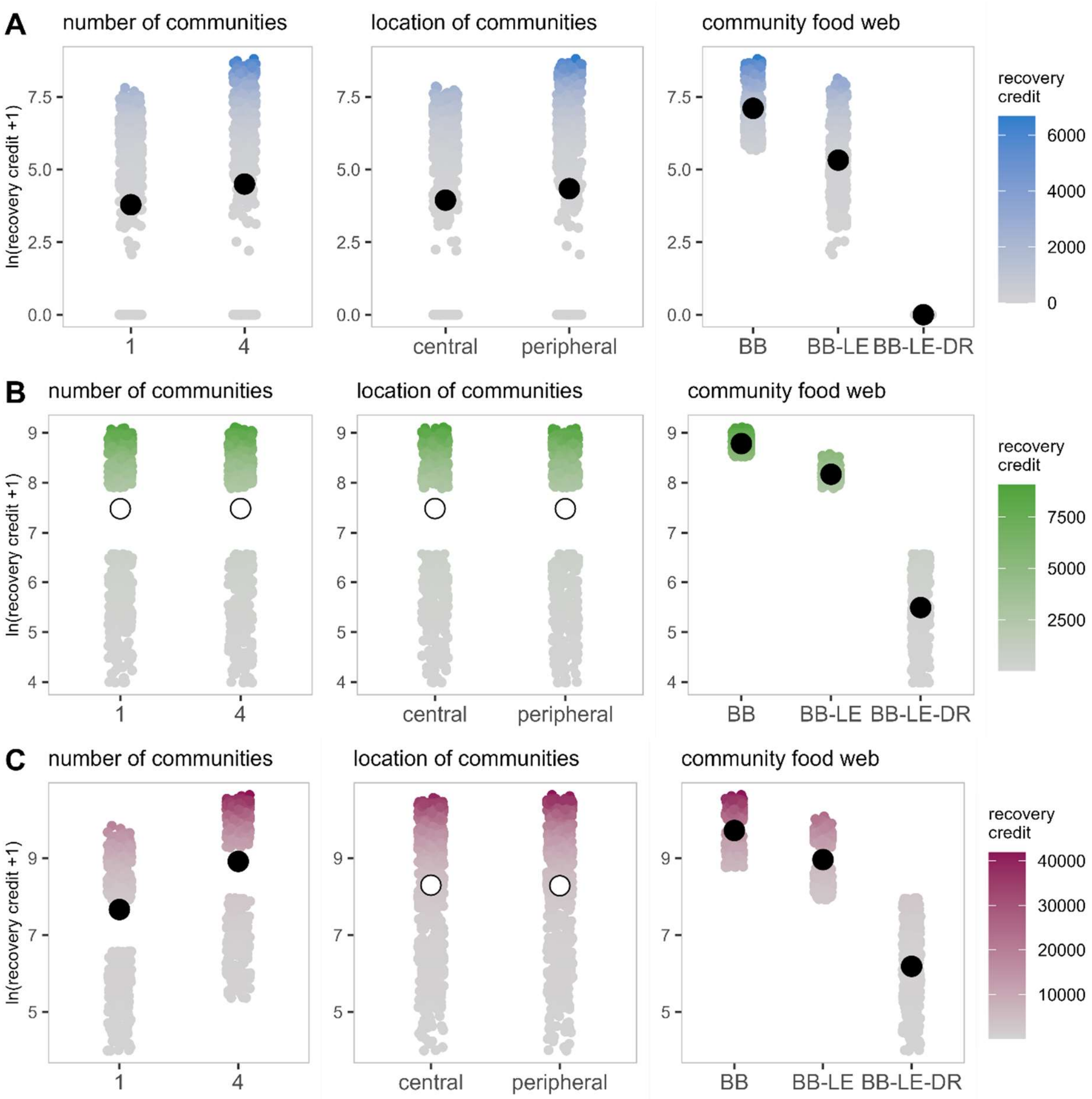


Figure S 11 Simulated recovery of aphid BB (A) in initially *empty* patches, (B) in initially *populated* patches, and (C) metapopulation. Panels show the effects of initial number of communities, location of initial communities, and community food web on the recovery credit ($ln(x+1)$ transformed). Smaller points represent recovery credit calculated from model simulations, with colours indicating their untransformed values. Larger points and vertical lines depict average linear model predictions and their 95% confidence intervals, respectively. Full and empty points indicate statistically significant and insignificant effects, respectively.

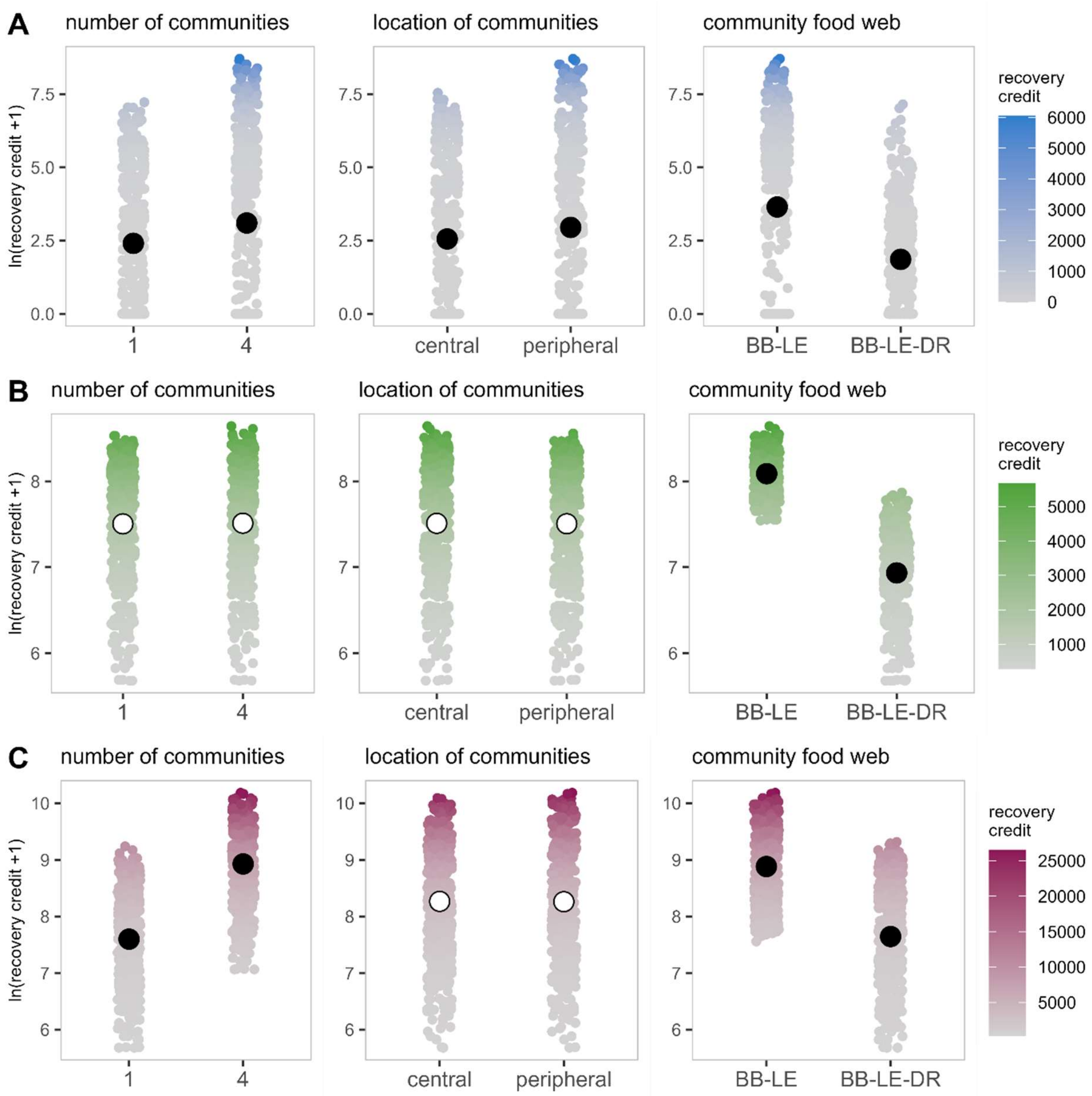


Figure S 12 Simulated recovery of aphid LE (A) in initially *empty* patches, (B) in initially *populated* patches, and (C) metapopulation. Panels show the effects of initial number of communities, location of initial communities, and community food web on the recovery credit ($ln(x + 1)$ transformed). Smaller points represent recovery credit calculated from model simulations, with colours indicating their untransformed values. Larger points and vertical lines depict average linear model predictions and their 95% confidence intervals, respectively. Full and empty points indicate statistically significant and insignificant effects, respectively.

#### Contributions to population change

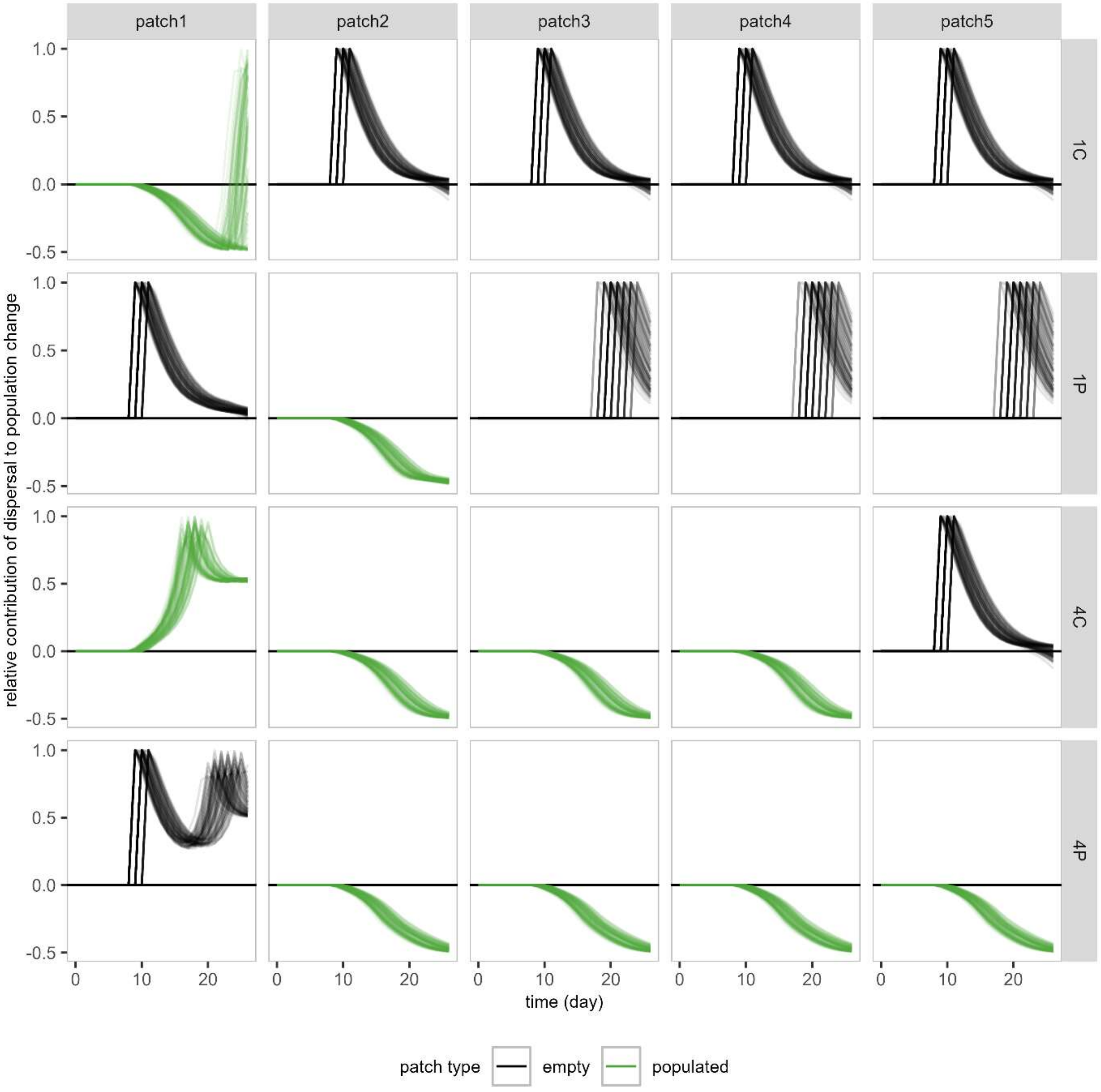


Figure S 13 Evolution of the relative contribution of dispersal to population change of aphid BB in *BB* community. Positive and negative contributions indicate net immigration and emigration, respectively. Panels correspond to patches (columns) in different landscapes (rows). Colours indicate the initial state of the patch. Lines join observations belonging to the same simulation replica.

### Simulated scenarios

#### Larger landscapes

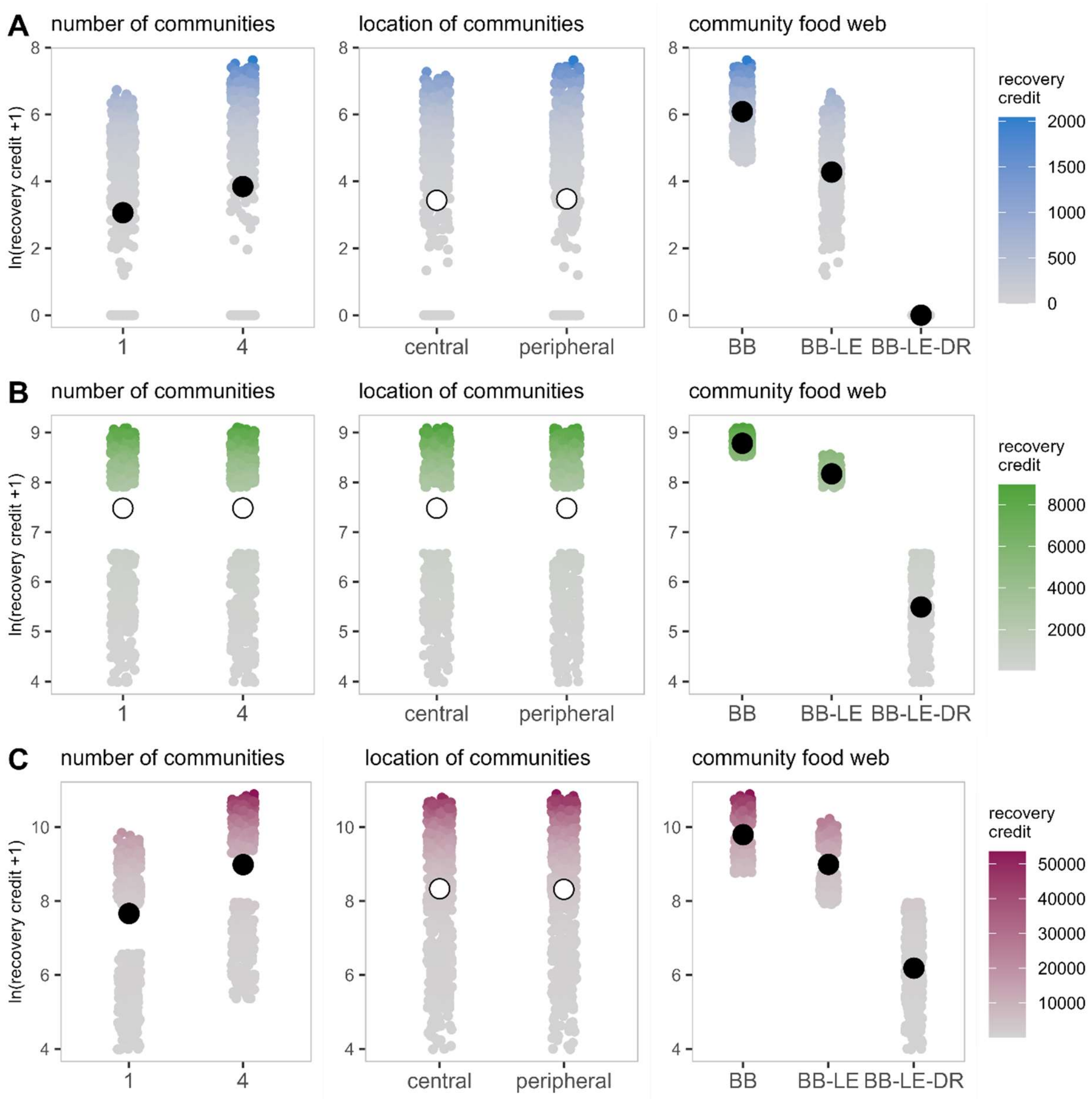


Figure S 14 Simulated recovery of aphid BB (A) in initially *empty* patches, (B) in initially *populated* patches, and (C) metapopulation in the *spread* patch configuration. Panels show the effects of initial number of communities, location of initial communities, and community food web on the recovery credit ($ln(x+1)$ transformed). Smaller points represent recovery credit calculated from model simulations, with colours indicating their untransformed values. Larger points and vertical lines depict average linear model predictions and their 95% confidence intervals, respectively. Full and empty points indicate statistically significant and insignificant effects, respectively.

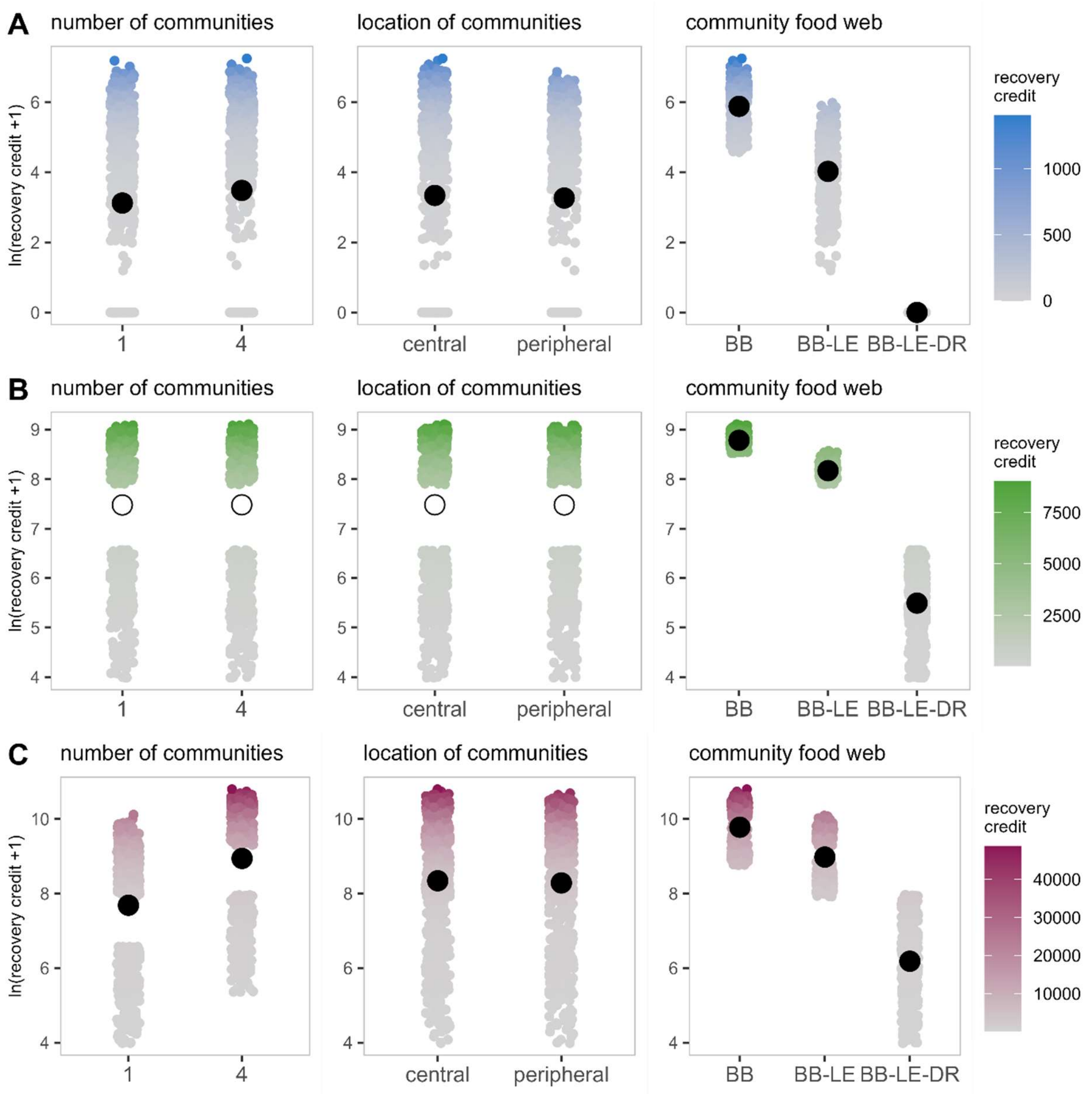


Figure S 15 Simulated recovery of aphid BB (A) in initially *empty* patches, (B) in initially *populated* patches, and (C) metapopulation in the *clustered* patch configuration. Panels show the effects of initial number of communities, location of initial communities, and community food web on the recovery credit ($ln(x + 1)$ transformed). Smaller points represent recovery credit calculated from model simulations, with colours indicating their untransformed values. Larger points and vertical lines depict average linear model predictions and their 95% confidence intervals, respectively. Full and empty points indicate statistically significant and insignificant effects, respectively.

#### Larger communities

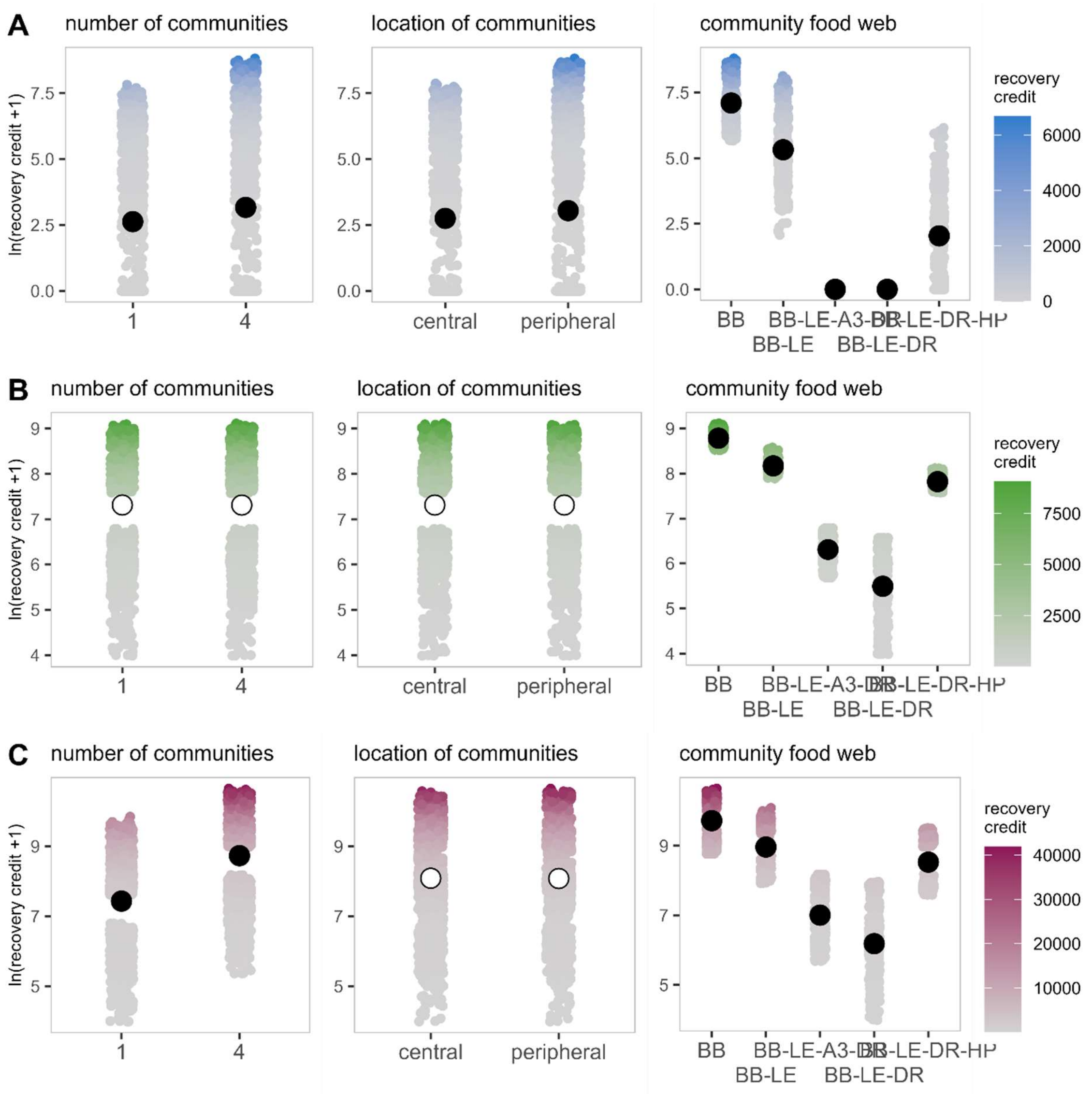


Figure S 16 Simulated recovery of aphid BB (A) in initially *empty* patches, (B) in initially *populated* patches, and (C) metapopulation, in simulations with larger communities. Panels show the effects of initial number of communities, location of initial communities, and community food web on the recovery credit ($ln(x + 1)$ transformed). Smaller points represent recovery credit calculated from model simulations, with colours indicating their untransformed values. Larger points and vertical lines depict average linear model predictions and their 95% confidence intervals, respectively. Full and empty points indicate statistically significant and insignificant effects, respectively.
